# Engineering Gelation Kinetics in Living Silk Hydrogels by Differential Dynamic Microscopy Microrheology and Machine Learning

**DOI:** 10.1101/2021.05.15.444303

**Authors:** Rhett L. Martineau, Alexandra V. Bayles, Chia-Suei Hung, Kristofer G. Reyes, Matthew E. Helgeson, Maneesh K. Gupta

## Abstract

Microbes embedded in hydrogels comprise one form of living material. Discovering formulations that balance potentially competing mechanical and biological properties in living hydrogels—for example gel time of the hydrogel formulation and viability of the embedded organisms—can be challenging. In this work, a pipeline is developed to automate characterization of the gel time of hydrogel formulations. Using this pipeline, living materials comprised of enzymatically crosslinked silk and embedded *E. coli*—formulated from within a 4D parameter space—are engineered to gel within a pre-selected timeframe. Gelation time is estimated using a novel adaptation of microrheology analysis using differential dynamic microscopy (DDM). In order to expedite the discovery of gelation regime boundaries, Bayesian machine learning models are deployed with optimal decision-making under uncertainty. The rate of learning is observed to vary between AI-assisted planning and human planning, with the fastest rate occurring during AI-assisted planning following a round of human planning. For a subset of formulations gelling within a targeted timeframe of 5-15 minutes, fluorophore production within the embedded cells is substantially similar across treatments, evidencing that gel time can be tuned independent of other material properties—at least over a finite range—while maintaining biological activity.

## 1. Introduction

A new frontier in materials development exists in combining living components with traditional material matrices. When living cells, for example bacteria, are encapsulated within traditional material matrices, new and potentially useful properties can be conferred. For example, embedding bacteria with inherent sensitivity to environmental signals such as pH or the presence of heavy metals^[1]^ can impart environmental awareness to a material. Enhancements are not limited to competencies which natively exist in embedded organisms; embedded cells can be genetically reprogrammed to be sensitive to non-native signals, as in the case of *S. Cerevisiae* engineered to be responsive to the presence of fungal pathogens.^[2]^ When the embedded cells are also capable of responding to sensed signals, for example by secreting growth factors,^[3]^ enzymes,^[4]^ or functionalized protein fibers,^[4]^ then sensing and actuation circuits are in place to enable smart, evolvable, and living biomaterials.^[5]^

A conceptual bacteria-enabled material is shown in **Figure 1**. In this illustration, genetically engineered bacteria are embedded into a hydrogel matrix. The cells possess a desired environmental sensitivity and are programmed to produce and secrete protein upon environmental stimulus. The matrix is designed for specific applications with regard to material stiffness and moldability, optical transparency, permeability to diffusing molecules including water and secreted proteins, and cure time.

**Figure 1.**
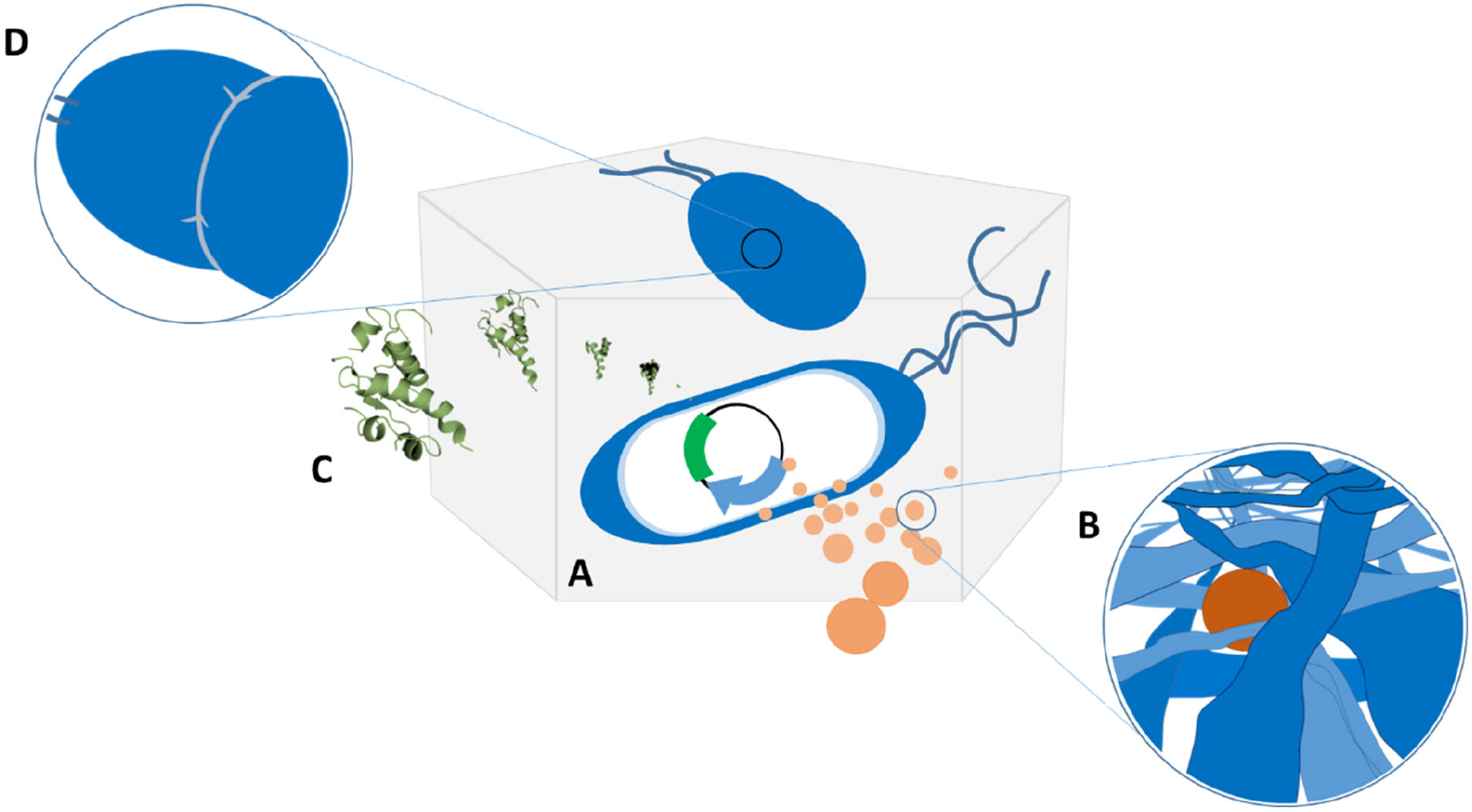
A generic bacteria-embedded living biomaterial. Living biomaterials can be formed by embedding microbes into a material matrix. To impart specific biological activity, the microbes may be genetically engineered (A). Signal molecules, (e.g. small molecules ∼1 nm in diameter), may be required to diffuse through the embedding medium to be internalized or otherwise sensed by the microbes (B). In response to signal molecules, the microbes may produce biological materials which are transported extracellularly and then diffuse through the embedding medium (C). For some embedding media, microbial mobility is limited and binary fission (D) is difficult, whereas for less stiff or more porous matrices, microbial mobility and reproduction are unhindered.

Care must be taken such that the formulation of the embedding matrix does not impede the intended biological activity. Chemical components in the formulation or resultant material properties may adversely impact the function of the biological components of the composite. Conversely, the presence or activity of the embedded microorganisms should not adversely impact the mechanical properties of the material. Discovering the formulations that balance the competing biological and materials requirements may be a difficult task, as not many tools exist for characterization of biomaterial formulations—especially the mechanical properties of those materials—in an automated or high-throughput fashion.

### 1.1 Living bacterial silk hydrogels

There is a significant body of literature describing efforts to embed living cells into hydrogels. Among hydrogels of recent interest, silk-based hydrogels are prominent.^[6]^ A number of properties make silk-based hydrogels attractive: in its natural, fibrous forms, silk has outstanding strength and elasticity.^[7]^ Silk is both biocompatible and biodegradable.^[6a, 6b, 6e–g, 8]^ When reconstituted from cocoons produced at industrial scale, silk is abundant. Furthermore, as a genetically encoded, proteinaceous material, a large number of varieties can potentially be produced using recombinant methods.^[9]^ In addition to sequence modifications at the genetic level, silk as a protein possesses handles for chemical functionalization and bioconjugation, further expanding the palette of material options for engineered applications.^[10]^ Finally, silk has been successfully processed from aqueous solutions into fibers, hydrogels, foams, and films.^[6e]^

Due to its favorable properties, silk materials have featured in numerous applications including: scaffolds for tissue engineering;^[6a, 8c, 11]^ injectable matrices for drug delivery;^[6d, 6g, 12]^ immobilization matrices for biocompatible, sub-dermal sensors;^[6d, 13]^ optical devices;^[8a, 14]^ and bio-functionalized bulk materials for microfluidic devices.^[15]^

Bacteria are an attractive option for adding biological function to materials due to their robustness and genetic tractability. Previous reports include embedding *Bacillus sphaericus* in hydrogels for calcium carbonate precipitation for self-healing concrete,^[16]^ *E. coli*^[17]^ and *B. subtilis*^[18]^ into 3D printing matrices for programmable secretion of biomolecules, *E. coli* into polyacrylamide and agarose for sustained secretion of drugs,^[19]^ *E. coli* into polyvinyl alcohol for heavy metal toxicity biosensing,^[1]^ and *Pseudomonas putida* and *A. xylinum* into hyaluronic acid, fumed silica, and κ-carrageenan for bioremediation and *in situ* production of bacterial nanocellulose.^[20]^

### 1.2 Linking formulation conditions with material properties

A key requirement for successfully designing living materials is linking formulation conditions with final material properties and—concurrently—biological activity.^[21]^ Material properties of general interest include those related to material formulation and processing (cure time, polymer fraction), mechanical properties (toughness, stiffness, fatigue strength), and transport properties (mass diffusivity, hydraulic permeability, etc.). Biological activity includes viability and motility of embedded organisms, ability to proliferate, responsivity of embedded organisms to signals of interest, or activity of secreted biologicals within or upon the embedding matrix. Generally speaking, trade-offs exist between idealized mechanical properties and idealized biological activity. A material’s suitability for a given application may depend critically upon finding an optimum among competing characteristics.

In developing a novel living material to achieve this balance of properties, a vast space of formula component permutations may need to be characterized. In the case of enzymatically crosslinked silk, hydrogels can be formed from aqueous solutions of purified silk protein, horseradish peroxidase (HRP) as an enzymatic crosslinker, and hydrogen peroxide.^[22]^ Gelation kinetics and final material stiffness vary widely over this multi-dimensional, continuous formulation space depending on the concentrations of the individual components.^[6e–h, 10–11, 22]^ The inclusion of microbes or other biological components within the embedding medium also impacts the material properties. Conversely, components in the embedding medium potentially impact the biological properties of the living material. As an illustrative example, Hasturk and coworkers embedded human mesenchymal stem cells into hydrogels comprised primarily of silk.^[6c]^ The materials were desired to rapidly gel once mixed from precursor solutions and to achieve both high stiffness and high bioactivity in the final hydrogel. Casting gels from precursors dispersed in distilled water resulted in poor survivability of the cells due to osmotic stress, but casting gels from precursors mixed into buffered salt solutions resulted in unacceptably slow gelation. To find formulation conditions that met all performance criteria, it was necessary to navigate a formulation space defined by a large number of parameters. Silk fraction, hydrogen peroxide concentration, crosslinking enzyme concentration, embedded cell concentration, buffer strength, and even the concentrations of potential crosslinking additives were all critical parameters impacting both the material and the biological properties of the final material. In the cited study, only a fraction of treatment combinations could be tested, possibly leaving the most suitable material formulation undiscovered.

### 1.3 Estimating gel times by automated DDM microrheology

Rapid screening of the mechanical properties of hydrogel formulations would facilitate living materials developments. For many applications, for example injectable living hydrogels with biological activity, the gel time of the material (i.e., the time scale over which solid-like mechanical properties develop) is one property that could critically impact a material’s successful deployment. Prior to this work, despite the existence of a number of techniques for gel time measurement in hydrogels (see **Supporting Information** sections **S1-S3**), a fully automatable process for gel time measurement in hydrogels had not been reported. Limitations of previous techniques impacting utility for automated implementation include large sample volumes, laborious setup, a need for user interactivity, and ambiguous gelation criteria.

Among existing techniques most promising for automated application is multiple particle tracking (MPT) microrheology^[23]^. Indeed, MPT has been used for high-throughput screening^[24]^. However, the major limitation of MPT precluding automated application is its need for user supervision, as the particle tracking algorithms generally require tuning of analysis parameters from sample to sample.

A new image analysis technique known as differential dynamic microscopy (DDM) has recently been adapted to conduct microrheological measurements.^[25]^ Like MPT microrheology, the technique requires commonly available optical equipment. In contrast with MPT microrheology, however, DDM requires no *a priori* input selection or interactive tuning^[25–26]^ and is therefore well-suited for automated application. Through Fourier analysis of image intensity fluctuations^[25a, 25c, 27]^ and the application of theories for thermally-driven motion, DDM quantifies the mean-squared displacement (MSD) of embedded tracer beads without the need to track individual particle trajectories.^[25b, 28]^ Once MSDs of gelling samples are obtained, a number of techniques can be applied to ascertain the gel time.^[24b, 24d, 29]^

### 1.4 Experimental approach and findings

In this work, we present a new experimental methodology and pipeline based on DDM to automate microrheology experiments for the determination of gel time. We also apply novel machine learning algorithms specifically formulated for characterizing a 4D formulation space in terms of gel time. As a use case, we develop a living silk hydrogel embedded with viable microorganisms, anticipating applications such as injectable and self-setting biological delivery materials. We asked the following coupled optimization questions: What bacteria-embedded silk hydrogel formulations produce gelation within 5-15 minutes? Within those formulations, which promote maximum bacterial activity?

In answer to these questions, we demonstrate and validate the capabilities of this new methodology through three experiments: 1) correlation of gel time estimation by a novel adaptation of DDM microrheology—requiring no user inputs and thus fully automatable—to gel time estimation by traditional methods based on MPT which do require user inputs, 2) characterization and prediction of gel time regime boundaries in bacteria-laden silk hydrogels spanning a 4D formulation space by use of novel machine learning algorithms and an automated gel time characterization pipeline, and 3) comparison of bacterial protein production in silk hydrogels of various formulations gelling in 5-15 minutes by use of confocal microscopy.

In brief, we find that DDM can be used as the basis to objectively and autonomously estimate gel times in silk hydrogels. Within a 4D formulation space, bacteria-embedded silk hydrogel formulations exhibited gel times ranging from instantaneous gelation to no gelation at all, with 8 of 63 treatments gelling within the targeted 5-15 minute window. Among a subset of samples that gelled within the 5-15 minute window, biological productivity of embedded *E. coli*, as measured by fluorescent protein production and colony morphology, was found to be relatively constant despite a significant difference in formula constituency.

Over the course of the 4D formula space characterization campaign, we employed Bayesian ML models and algorithms to perform decision-making under uncertainty to accelerate knowledge acquisition. ML supervision can proceed without operator intervention, however the fastest rate of learning was observed when a round of expert user decision making was interspersed between rounds of AI decision making. Without operator intervention, the ML algorithms were capable of approximating gel regime boundaries in as few as 20 treatments.

The findings of this work collectively illustrate the complex relationship between formulation and properties in living silk hydrogels and show that automated materials characterization techniques such as DDM microrheology can help to navigate seemingly intractable and vast formulation spaces, especially when paired with ML-informed modeling and decision making algorithms. Importantly, the techniques deployed in this work—in particular gel time estimation by DDM—are easily transferred to other laboratories and can be implemented on equipment which is readily available.

## 2. Results and Discussion

### 2.1 Comparison of gel time estimation by DDM and by MPT

Characterization of gelation kinetics requires an estimation of the gel time. A number of methods for gel time estimation exist based on various dynamic linear viscoelastic material functions, and these methods are summarized in detail in the **Supporting Information Sections S1-S3**. These methods include determination of the point of crossover between the elastic and viscous moduli (*G′* ∼ *G′′*) for a given deformation frequency or frequency range, the point at which the loss tangent tan *δ* becomes independent of frequency, or by time-cure superposition, in which time-domain shifting of the viscoelastic response is used to identify the asymptotic point at which shift factors transition from pre-gel to post-gel scaling. Each of these methods is accessible to MPT microrheology by way of MSD curves. However, MPT microrheology presents unique difficulties and limitations for automated determination of gel points, either due to a lack of theoretical support for precisely defining the gelation point, or due to user inputs or interventions that are required to ensure robustness of the method.

As mentioned previously, DDM can be used to produce MSDs without user supervision and is thus potentially amenable to automation. When used to calculate MSD curves, DDM compares well with MPT over a wide variety of conditions.^[25b, 28]^ As MSD curves can serve as the foundation for rigorous microrheological assessment of gelation, both DDM and MPT can be used to measure gel time.

During a gelation reaction, probe particles undergo progressively smaller cumulative displacements approaching the gel point and beyond, where probe particle motion is restricted by the elasticity of the gel. DDM fails to replicate MPT-extracted MSD curves in samples exhibiting very small cumulative displacements (see **Supporting Information Section S4**), where the weak intensity fluctuations associated with particle motion are not easily discernable from other contributors to the image structure function.^[25b]^ Under these circumstances, the DDM signal generally fails to meet the tolerance criterion for successful MSD estimation. In some cases under these circumstances, the tolerance criterion is met but no physically meaningful MSD curves are produced. An example of the difference in the sensitivity range between MPT and DDM is highlighted in **Section S4** of the **Supporting Information**, where MPT yields physically reasonable MSD curves at time points well beyond those that produce physically reasonable MSD curves by DDM.

The lack of MSD curves resulting from DDM analyses of late-stage gelation limits the amount of data that can be observed during a gelation reaction, potentially precluding gel time estimation by *G*′/*G*′′ cross-over, time-cure superposition, or tan(*δ*) analysis. However, in practice, the point at which DDM fails to extract physically reasonable MSD curves may itself be useful as an estimate of the gel time. In this work, we develop a technique for gel time estimation based on identifying the time during a gelation reaction after which no MSD curves can be extracted from *D*(*q, Δt*) curves obtained by DDM. We term this method DDM-RL, referring to the resolution limit of DDM for MSD extraction. If DDM-RL does in fact produce useful estimates of the gel time, the advantages of DDM over MPT—namely, the parameter-free nature of the analysis—make DDM microrheology an excellent basis for an automatable pipeline for objective gel time characterization.

The first investigation presented in this work examines DDM-RL as a method for gel time estimation. Estimates obtained by DDM-RL are compared with estimates obtained by converting MSDs obtained from MPT into mechanical moduli and then observing the time at which *G*′ exceeds *G*′′ over all measured frequencies (termed here, MPT-*G*′/*G*′′).

Passive probe microrheology data from three silk hydrogel formulations with distinct gel times were analyzed by both DDM-RL and MPT-*G*′/*G*′′. For this comparison, none of the silk formulas contained bacterial biomass. All gels contained 3% silk (w/v). The HRP and H_2_O_2_ concentrations varied as: 1) 20 U/ml HRP with 53 ppm H_2_O_2_, 2) 10 U/ml with 25 ppm H_2_O_2_, and 3) 3.33 U/ml HRP with 25 ppm H_2_O_2_. Results are shown in **Figure 2**. Estimates by DDM-RL are reported as the sum of ½ the sampling interval and the time associated with the last physically reasonable MSD curve obtained by DDM. Errors are reported as ± ½ the sampling interval. See **Section 4** for experimental detail. Estimates by MPT-*G*′/*G*′′ are reported as the range of times bracketing the presumed switch from a pre-gel to post-gel regime based on the criterion that *G*′ > *G*′′ (or *G*′ ∼ *G*′′) for all measured frequencies.

**Figure 2.**
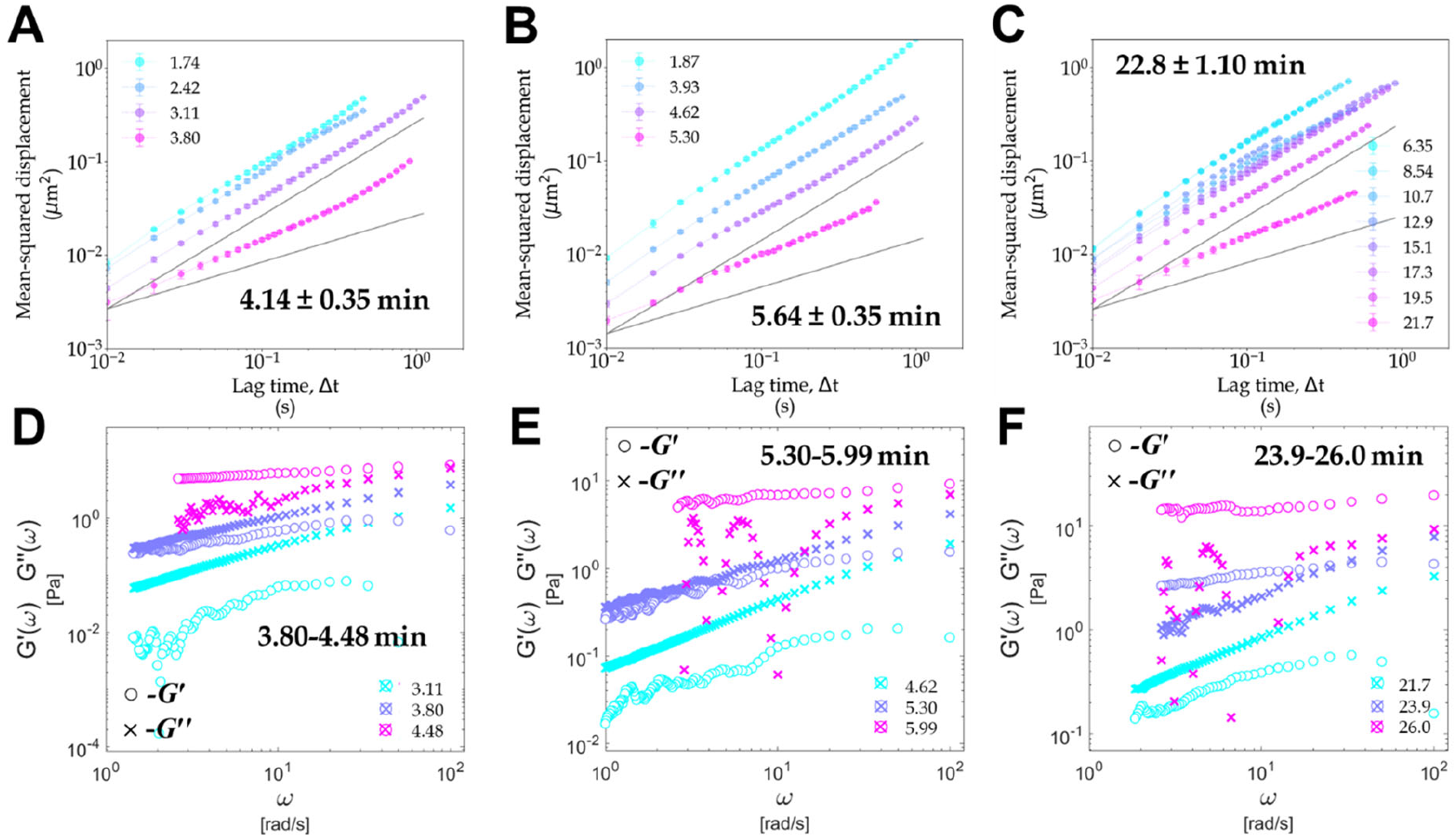
Estimation of gel time by DDM-RL and MPT-*G*′/*G*′′. Gel time was estimated by DDM-RL (Panels A-C) and by MPT-*G*′/*G*′′ crossover (Panels D-F) for three gels. Each gel contained 3% silk but varied in the other constituents. For the sample comprised of 20 U/ml HRP with 53 ppm H_2_O_2_, gel times were measured at 4.14 ± 0.35 minutes by DDM-RL (A) and between 3.80 and 4.48 minutes by MPT-*G*′/*G*′′ (D). For the sample comprised of 10 U/ml HRP with 25 ppm H_2_O_2_, estimates were 5.64 ± 0.35 minutes by DDM-RL (B) and between 5.30 and 5.99 minutes by MPT-*G*′/*G*′′ (E). The sample comprised of 3.33 U/ml HRP with 25 ppm H_2_O_2_, was measured to gel at 22.8 ± 1.10 minutes by DDM-RL (C) and between 23.9 and 26.0 minutes by MPT-*G*′/*G*′′ (F). Grey lines indicate power law slopes of ½ and 1. Panels D-F show *G*′ and *G*′′ plots obtained by local power-law fitting of MSDs obtained by MPT.

In general, during gelation processes, the MSDs measured by both DDM and MPT are expected to exhibit power law behavior with decreasing power law exponents as gelation proceeds, i.e., a decreasing linear slope on a log-log scale. For the cases of DDM here, each sample exhibits power law slopes greater than 0.5 for all time points at which the MSD can be resolved. After converting MSDs obtained by DDM to *G*′/*G*′′ plots, no time point exhibited *G*′ in excess of *G*′′ for all frequencies. This is due to the resolution limit of DDM in extracting MSDs, with a lower bound on MSD related to the pixel size and the influence of intensity fluctuations away from the focal plane.^[25b]^ However, DDM does produce MSDs for some time points where *G*′> *G*′′ for some frequencies. In the case of 3% silk, 10 U/ml HRP with 25 ppm H_2_O_2_ (**Figure 2B**), DDM produced an MSD curve for the 5.3 minute time point that corresponds to *G*′∼*G*′′ for many of the lower frequencies. This demonstrates that DDM can resolve MSD curves at time points very close to the gel time. This observation also highlights that the number of observations taken during a gelation reaction is an important parameter in accurately measuring gel time by DDM-RL. Specifically, maximizing the number of video stacks taken during a gelation reaction—without compromising the frequency content of those video stacks—is important for achieving the most precise estimates of the gel time by DDM-RL.

In the three cases presented in **Figure 2**, estimates of gel time by DDM-RL and by MPT-*G*′/*G*′′ approximately agree. Likewise, for an additional three replicates of the 3% silk, 3.33 U/ml HRP, and 25 ppm H_2_O_2_ case, each estimate by DDM-RL overlaps with the estimate by MPT-*G*′/*G*′′ (data not shown). In most cases, the latest gel time estimated by DDM-RL matches the earliest time suggested by MPT-*G*′/*G*′′. That is, DDM typically fails to resolve MSDs when *G*′ begins to be equal to or to exceed *G*′′ for some frequencies. DDM-RL therefore tends to slightly underestimate the gel time. It is also possible that DDM-RL reports gelation when gelation has not (and will not) occur, although this hasn’t been observed in any of the cases reported in this study. On the other hand, if drift and convective currents exist in the sample (as might be indicated in Fig2A at the 3.8 minute time point, for example), DDM-RL might tend to overestimate the gel time. Indeed, many of the MSD curves reported in this study exhibit upturns in the MSDs at large lag times. Dedrifting algorithms applied to the MPT data eliminate these upturns (See **Supporting Information S5**). The drift is not unexpected as the measurements were taken in open well dishes to facilitate robotic handling. Importantly, the drift in these samples does not appear to significantly impact the gel time estimates by DDM-RL.

In its current formulation, DDM does not identify drift or convective currents in the sample. A potentially useful adaptation to the DDM algorithm could be developed to identify and perhaps computationally eliminate excessive drift or convective currents, as is currently possible with MPT.

Despite the difficulties of gel time estimation by DDM, DDM still appears to be useful in screening scenarios where automated analysis is desirable. Each of the samples observed to gel by way of MPT-*G*′/*G*′′ crossover was also successfully reported to gel by DDM-RL, with estimates of gel time generally agreeing within one measurement interval.

Based on these findings, we argue that for objective, parameter-free characterization of gel time in hydrogel formulations, DDM-RL produces approximate gel times that are sufficiently accurate to be useful as an initial screen. Although the estimation of gel time by this technique is not rigorously supported by theory, the MSD curves collected during execution of the technique do capture important rheological information that is quantitatively comparable to more rigorous viscoelastic measures obtained by MPT. As desired, the MSDs obtained by DDM can be processed to quantify the evolution of complex shear modulus or viscosity as a sample gels. Also, since the same raw data is used for both DDM and MPT, the image stacks for a given sample can be reprocessed by MPT if additional information is required—for example the explicit particle trajectories or van Hove displacement distributions.

As already mentioned, DDM does not require a user to fine-tune parameters for each analysis and is therefore an excellent option for automated characterizations. This is especially beneficial when the sample is changing over the course of an experiment, as was sometimes observed for these gelling samples. As samples gelled, changes in the opacity required careful tuning of the processing parameters when analyzing by MPT. In contrast, analysis by DDM proceeded without regard for changes in optical opacity.

A typical gel time estimation experiment by DDM-RL took approximately 1.5 hours to complete. Each formula treatment to be analyzed for gel time was observed microscopically for up to 30 minutes and then processed by DDM for the next approximately 45 minutes. A typical data set consists of up to 15 image stacks of 1000 images each, with each stack requiring about 5 minutes to process when the images were processed using exponential lag time spacing. In contrast, processing by MPT required approximately 30-45 minutes per image stack, with intensive user interaction for the first 5-10 minutes. Once gel times were determined, results were supplied to the ML experimental planner which required approximately 10 minutes to produce recommendations for the next iteration’s formulas.

Parameter-free measurement of evolving MSD curves in gelling samples could have potentially been obtained using a recently developed convolutional neural network (CNN).^[30]^ The CNN was trained using both simulated and real particle imaging videos and can convert image stacks into MSDs without user-supplied parameters. We processed image stacks from a silk gelation (see aitracker.net) and observed that CNN based particle tracking and conversion to MSDs was excellent—and agreed well with DDM-based MSDs—for early time points. However, for later stages in the gelation process, the CNN algorithms failed. See **Supporting Information S6** for details which indicate that automated gel time estimation by CNN holds significant promise but is not currently suitable for routine use.

### 2.2 Automated screening of gel time in bacteria-laden silk hydrogels in 4D formulation space

Having established that DDM-RL is a useful estimator of gel time, we constructed a pipeline for its automated implementation. The pipeline consists of the following operations (**Figure 3A**): 1) formula specification, 2) mixing and dispensing, 3) video microscopy, 4) DDM microrheology, and 5) gel time estimation by DDM-RL. Once the gel time was determined, the experiment’s results were added to a growing ensemble of results, and a new formula specification was generated by an experimental planner and the cycle repeated.

**Figure 3.**
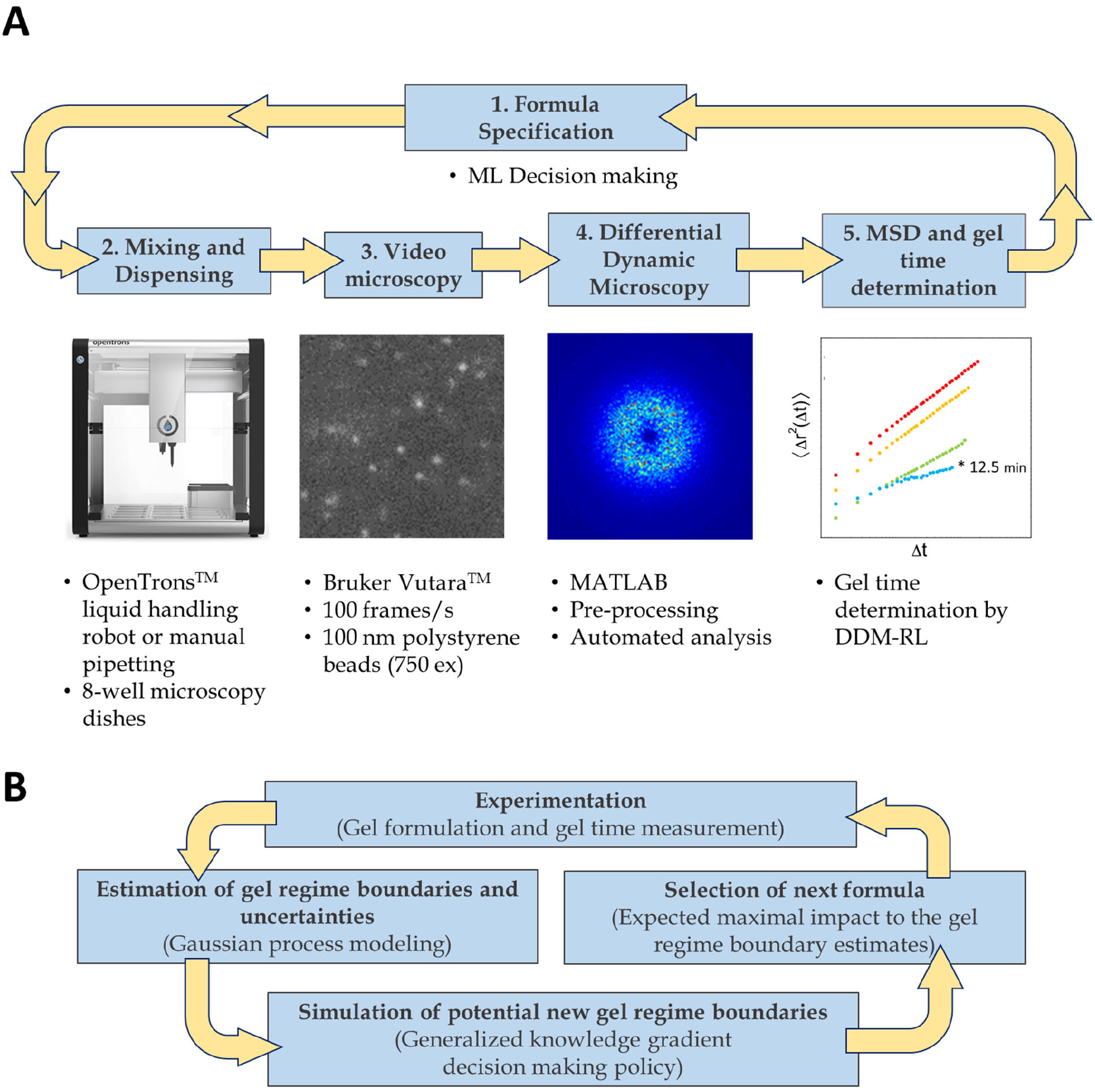
Automated gel time characterization pipeline. A closed-loop experimental campaign to characterize a 4D silk hydrogel formulation space was conducted iteratively (**Panel A**). Each iterate of the campaign began with a formulation specification suggested by a decision-making policy informed by Bayesian ML models of gelation regimes, or by an expert operator. Next, the suggested formulas were mixed using a pipetting robot or manually.

Samples were loaded into open wells and manually placed onto the microscope for image acquisition. Images were acquired automatically at set intervals and framerates and then processed using a series of automated MATLAB scripts which implemented DDM. The gel time was estimated from the time course of MSD curves using DDM-RL. Specific steps in the ML decision making policy are broken out in **Panel B** and described in the following.

The possible combinations of formula components in a 4D, continuous formulation space are potentially overwhelming. To address this issue, we deployed a novel ML strategy based on iterative Bayesian closed-loop design. In contrast to more traditional design of experiment (DOE) approaches based on collecting batches of observations to fill a model design space, we coupled small sets of exploratory experiments with Bayesian beliefs to more adaptively navigate the formulation space, relying on a sequential experimental design and a decision policy known as the Generalized Knowledge Gradient (GKG). The method was specifically devised to optimally identify gel regime boundaries in the design space. The basic scheme is illustrated in **Figure 3B**. After an experiment, the gel time landscape over the 4D input space is expressed as a Gaussian process (GP) model and used to estimate the gel regime boundaries. The GKG policy then selects the formulation that is expected to most impactfully change this prior estimate of the boundary. The policy employs a Monte Carlo (MC) approach to calculate the expected change of the boundary. Namely, the GP model is then also used to simulate new data points within the gelation time landscape from which new potential posterior gel regime boundaries are calculated and compared with the prior boundaries, obtaining a measure of difference between the two estimated boundaries. This is repeated several times to obtain an average measure of difference for a given formulation. The formula specification which is expected to maximally impact the gel regime boundary, as measured by this MC-averaged difference, is then selected as the next physical experiment to run, and the cycle repeats.

By focusing on specific desiderata in an adaptive manner, we can drive the exploration to achieve the particular design objectives of an investigation—in this case, finding formulas that gel within 5 to 15 minutes—without needing to learn the design space landscape in general. As a result, fewer experiments may be required, as has been demonstrated for a number of related sequential approaches in comparison with traditional global or space filling designs.^[31]^ This efficiency is particularly important in higher-dimensional design spaces as is the case in this application. The specifics of the boundary-learning ML method we developed for this work are presented in **Section 4-Experimental**, and in **Section S7** of the **Supporting Information**.

The results of a campaign to characterize gel times by DDM-RL in a 4D formulation space are presented below in **Figure 4**. The results were obtained from 63 formula characterization experiments. The formula samples were organized into three gel regimes: samples gelling within the selected 5-15 minute window, samples gelling faster than 5 minutes, and samples taking longer than 15 minutes to gel or not gelling at all. Individual observations are shown in the plots in **Figure 4A** which also show predicted gel regime boundaries as the campaign progresses.

**Figure 4.**
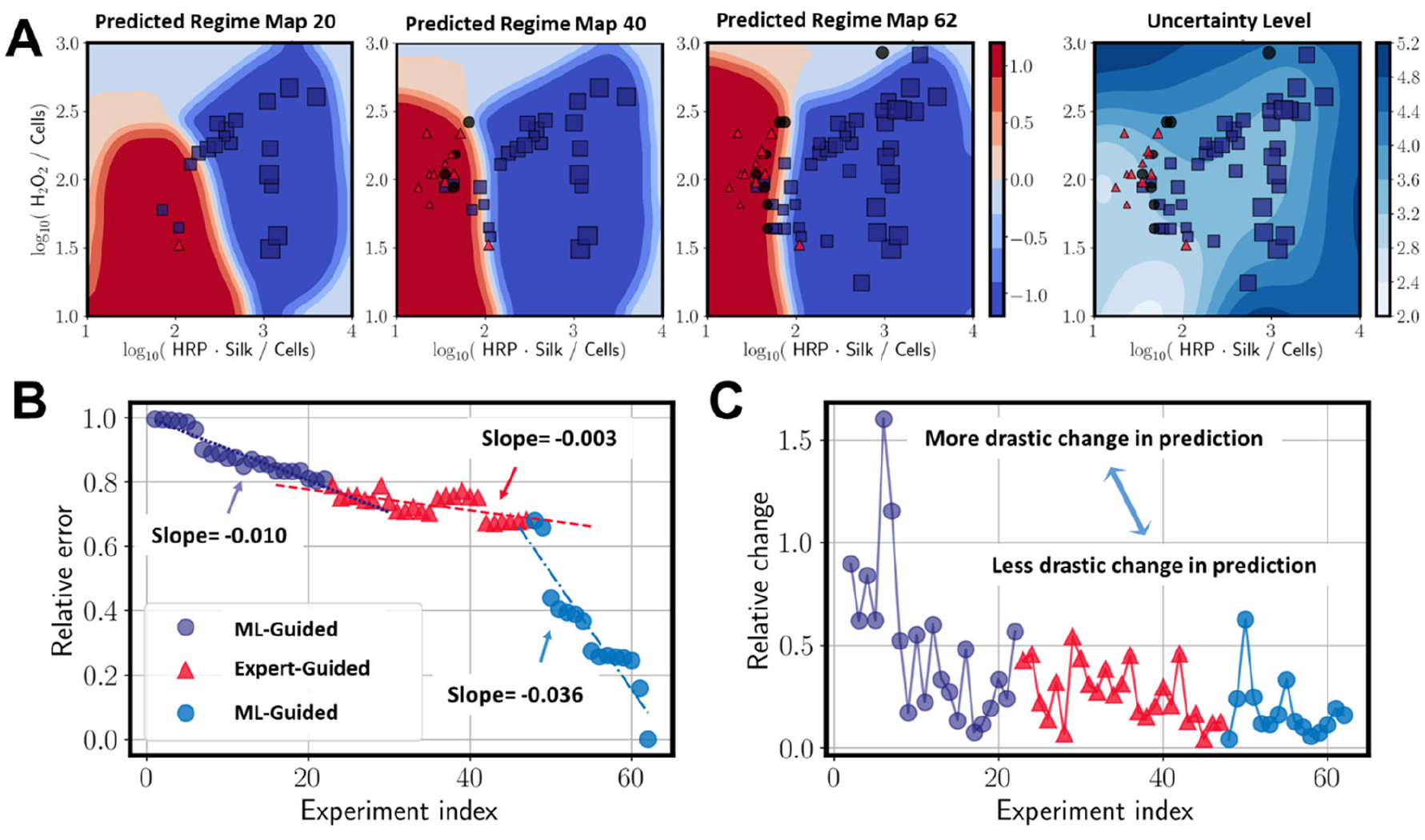
ML and expert-guided experiment planning. **Panel A** presents low-dimensional visualizations of the mean-estimates of the gelation regimes after 20, 40 and 62 experiments. Observations that gelled within the targeted window of 5-15 minutes are shown as black circles. Observations that gelled faster than 5 minutes are shown as blue squares. Observations that took longer than 15 minutes, or that didn’t gel at all in the 30 minute observation window, are marked as red triangles. The size of the marker reflects the relative concentration of HRP, with the larger markers indicating larger concentrations. Colored regions indicate areas in the formulation space predicted by an SVM classifier to produce samples gelling in three gelation regimes: 1) faster than 5 minutes (blue region), slower than 15 minutes or not gelling at all within the observation period of 30 minutes (red region), or 3) samples gelling within the targeted window of 5-15 minutes (lighter red and blue). Units are coded gel regime units, with **l**=-1 representing gel times less than 5 minutes and **l**=1 representing gel times greater than 15. The plot at far right in **Panel A** presents the uncertainty associated with the predictions after 62 experiments, representing the error bars (in minutes) on the corresponding gel regime predictions. **Panel B** plots the error in the estimates of the full-dimensional gelation time boundary obtained for each iteration in the campaign relative to the estimate obtained after 63 observations. Purple dots show the relative error during an initial ML-guided phase of the campaign. Red triangles show the errors observed during a phase in which a domain-expert selected the formula components with the intent to find samples that gelled within the desired timeframe. Blue circles show errors incurred during a second ML-guided phase of the campaign. Dashed lines are a linear fit to the data, with the slopes of these lines indicating a measure of the rate of learning, with more negative rates signifying faster learning. The expert opinion phase had the slowest rate of 0.003, which is more than 3 times slower than that of the initial ML-guided phase (rate = 0.01) and ten times slower than the final ML-guided phase (rate = 0.036). **Panel C** shows a comparison of successive gel time regime boundary estimates, as measured by relative change. Larger values indicate experiments that more drastically change the prediction. The most drastic changes occur in the beginning of the first ML-guided phase of the campaign.

Selection of formulations for characterization was alternated between our machine learning algorithm (picking four treatments per iteration) and a domain expert (picking one treatment at a time) attempting to pick treatment combinations that gelled within the 5-15 minute window based on domain expertise.

Gelation data and the Bayesian beliefs on gelation time, as presented in **Figure 4**, have been projected from full 4D input space to two dimensions by plotting as H_2_O_2_/Cell vs. HRP*Silk/Cells. Exploratory data visualization revealed this representation to be the most differentiating with respect to gelation kinetics.

Not all observations fell within regions delineated by the model predictions. Notably, though, the approximate gel regime boundaries are qualitatively projected in as little as 20 observations, with only minor alterations observed progressing through the full 63 experiments. Interestingly, the learning rate observed at the beginning of the experimental campaign is substantially slower than that observed in the later stages of the campaign. In both ML-guided phases, learning was faster than that achieved when a domain expert attempted to find regions within the targeted 5-15 minute window, suggesting the policy designed specifically to quickly learn the gel regime boundaries is successfully doing so. Of additional note, the treatments selected during the expert-guided stage appeared to result in an increased learning rate in the following ML-guided stage. Experiments selected by the expert were exploitation ones – experiments believed by the domain expert to produce gelation times in the target window, while ML-selected experiments attempted to balance exploitation and parameter space exploration. These results suggest alternative exploration/exploitation stages may be beneficial. Another interesting question raised here is the potential for collaborative teaming between ML algorithms and domain experts. The use of ML allows for reasoning in high dimensions with statistical analysis, while human experts can make suggestions using prior or unmodeled knowledge not built into the ML models. These results point to the beneficial effect that combining both approaches has to accelerating discovery.

The rapid rate of learning at the end of the 63 experiments suggests that more data are necessary to definitively characterize the gel regime boundaries. However, the decreasing error between each iteration’s expectations and the expectations in the final iteration suggests that predictive quality in the dataset is converging.

The gelation times of silk hydrogels in this study varied widely from near instantaneous gelation, to gelation over the course of tens of minutes, to not gelling at all even after many hours (as observed incidentally; the observation period for microrheology was 30 minutes). The formulation transition from samples that gel faster than 5 minutes to samples that take longer than 15 minutes to gel is abrupt; the gelation window of 5-15 minutes is extremely narrow under the conditions explored in this study. This is an example of a situation where discovering the gel regime boundaries may be preferable to fully characterizing the formulation space, as a full characterization is expensive to obtain and a sparsely occupied space-filling design might poorly characterize the boundaries.

Also indicated in **Figure 4C** is the relative concentration of HRP in each formulation. The size of the marker indicates the relative concentration of HRP, with larger markers indicating higher concentrations of HRP. A clear relationship exists between HRP concentration and gel time, with higher HRP concentrations tending to gel the reactions more quickly. This finding agrees with other literature detailing HRP-mediated silk hydrogelation.^[6c]^

**Table S1** (see **Supporting Information**) presents the data in tabular form. Silk was added in the range of 1.71-6.0% w/w, hydrogen peroxide ranged from 15-300 parts per million, HRP from 5-243 U/ml, and cell concentrations from 0.1 – 1.4 x 10^9^ cells/ml. 8 of the 63 samples, 7 obtained during the expert guided phase, gelled within the specified 5-15 minute window. Note that the object of the ML algorithm was to discover the gel regime boundaries in the formulation space and not to find specific formulas that gelled within the 5-15 minute window.

### 2.3 Comparison of protein production in 5-15 minute silk hydrogels

A second experiment was designed to determine which formulas that gelled within 5-15 minutes promoted maximum bacterial activity. Four such formulas were selected for further study. To facilitate comparison, each chosen formula contained a matched bacterial load of 1.14 x 10^9^ cells/ml. Beyond that, the selected formulas sampled the widest range of constituent concentrations available in the previously obtained 5-15 minute samples and included: 1) 5.31% silk, 175 ppm H_2_O_2_, and 10 U/ml HRP, 2) 3% silk, 50 ppm H_2_O_2_, and 18 U/ml HRP, 3) 2% silk, 125 ppm H_2_O_2_, and 20 U/ml HRP, and 4) 3% silk, 300 ppm H_2_O_2_, and 28 U/ml HRP.

For these experiments, fresh cultures were grown to the required OD_600_ and added to the treatments immediately. This is in contrast to the way cells were prepared and added to the treatments in the 4D formula space experiment presented in **Figure 4.** In that case, all cells for the entire campaign were prepared in one large batch and frozen in aliquots to ensure matched cell fractions throughout the characterization campaign. In these experiments addressing bacterial viability, fresh cells were added to gel formulas. Once fresh cells were added to the formulas, the samples were allowed to gel, overlaid with culture medium, and cultured for 24 hours under inducing conditions.

From preliminary growth experiments, it had been observed that both the fluorescence intensity produced by the embedded cells and evidence of extended bacterial growth (as indicated by the appearance of bacterial colonies) varied significantly based on location within a gel. That is, the further away a cell was from the nutrient overlay within any given sample, the less intense, generally, the fluorescence. In fact, induction of fluorescence in cells located deep within a gel was extremely poor (see **Section S8** of the **Supporting Information**.) In gels seeded with dilute suspensions of cells, where the oxygen and other nutrient levels were not limiting during early stages of growth, a number of distinct colony morphologies (including oblong spherical, filamentous, and ‘hairy’ filamentous) were produced in different hydrogel formulations even in deep sections (see **Section S9** in the **Supporting Information**). In contrast, when cells were added in higher concentrations (such as was done for the replicated treatments), cells grew into colonies only near the surface of the gel-liquid interface perhaps as facilitated by more readily available oxygen or other nutrients.

To ensure similar nutrient and oxygen availability between samples, gels were cast into thin sheets by adding approximately 100 μl of pre-gel material to the open wells and then spreading the material to cover the entire well bottom (∼ 1 cm^2^). Casting into thin sheets created areas that could be observed microscopically to contain cells growing in gels near the liquid-gel interface, where a pathway to oxygen and nutrient exchange was not blocked by competing cells. Confocal stacks used for comparisons between treatments were selected to show regions spanning the coverglass, at various z-positions within the hydrogel including near to the gel-liquid interface, and extending into the culture medium overlay. Thus each imaged gel section was no thicker than approximately 180 μm. **Figure 5A** illustrates the imaged regions. Results from imaging through the thin sections of the four bacteria-laden silk hydrogels cast in formulas that previously gelled within 5-15 minutes is presented in **Figure 5**.

**Figure 5.**
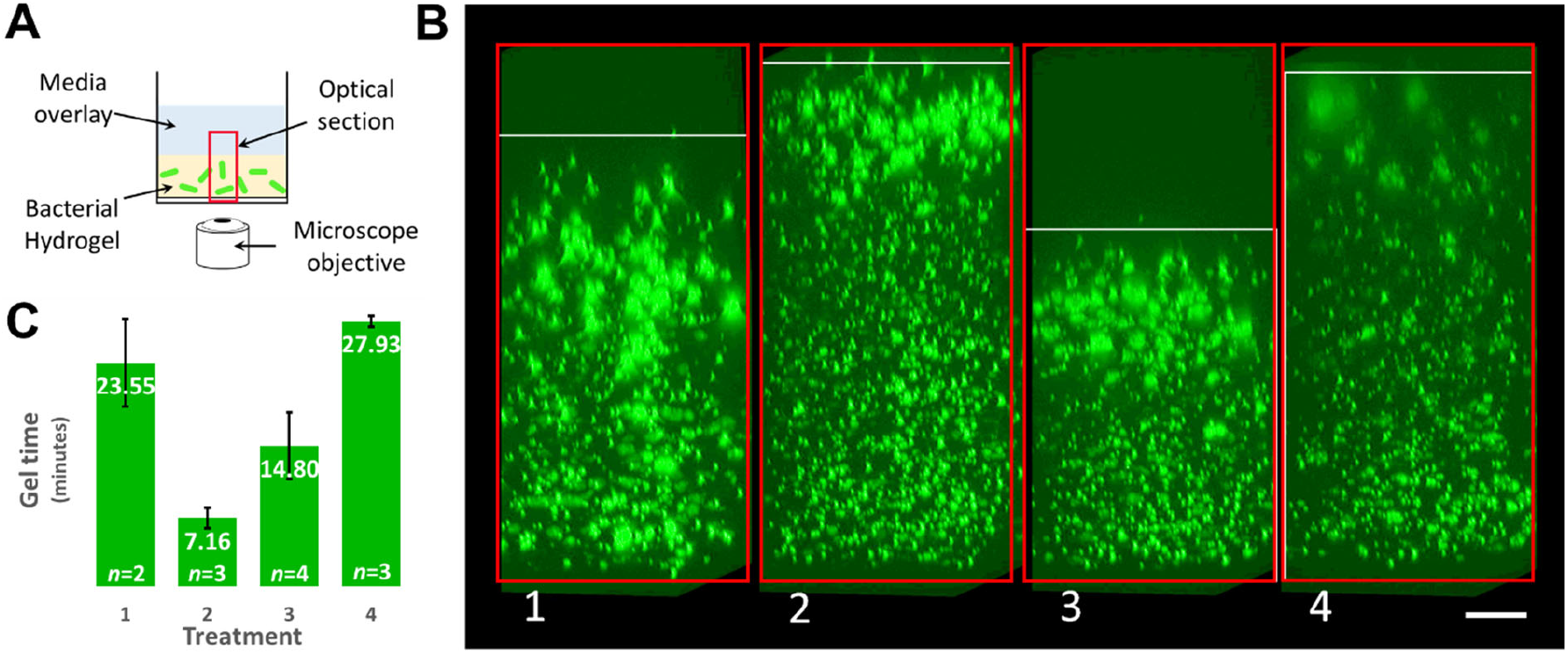
Overnight protein production in 5-15 minute silk hydrogel formulas. **Panel A** illustrates the confocal imaging setup. A relatively thin section of bacteria-laden hydrogel was imaged in an inverted configuration, with images spanning the region beneath the coverglass, through the hydrogel sample, and into the overlaying culture medium. **Panel B** shows representative confocal micrographs of bacterial cells growing near the gel-medium interface for four treatments that were previously observed to gel within 5-15 minutes. Each confocal image is a side view of the maximum intensity projection of a stack of 80 x 80 μm^2^ images sampled every 0.5 μm over a depth of 200 μm. Micrograph orientation matches the orientation depicted in **Panel A**. Cells positioned toward the top side of the images were progressively closer to the gel-medium interface, and gel sections were chosen such that the top side of each image shows (as a lack of fluorescence) the liquid media overlaying the hydrogel. The white rectangles mark the estimated location of the gelled material, whereas red rectangles illustrate the full stack of images taken through the sample. **Panel C** shows the gel times for each replicated treatment measured by DDM-RL, with gel times reported in minutes and the number of replicates for each treatment indicated. Treatment formulas included 1) 5.31% silk, 175 ppm H_2_O_2_, and 10 U/ml HRP, 2) 3% silk, 50 ppm H_2_O_2_, and 18 U/ml HRP, 3) 2% silk, 125 ppm H_2_O_2_, and 20 U/ml HRP, and 4) 3% silk, 300 ppm H_2_O_2_, and 28 U/ml HRP. Scale bar is 20 μm.

Of the four treatments recast with fresh cells, two produced gels that consistently gelled in the expected 5-15 minute window. These treatments were: 3% silk, 50 ppm H_2_O_2_, and 18 U/ml HRP; and 2% silk, 125 ppm H_2_O_2_, and 20 U/ml HRP. These growth experiments, with concurrent gel time measurements, were repeated on two, three, and four separate days (see **Figure 5B**) starting from fresh inoculum. In general, the repeatability of each treatment across days is excellent.

The two treatments that did not gel within the expected window (treatments 1 and 4 from **Figure 5B**) were observed to gel within 30 minutes. The inability to repeat gel times in all treatments of the growth experiments compared with the gel times observed for those matched treatments in the 4D characterization campaign may be due to a number of factors including the difference in cell seeding stock (fresh vs. frozen), variability in the age or activity of the other hydrogel precursors, or the discrepancy in reaction temperature between the gel time screening campaign and the campaign to study bioactivity in the matched subset of formulas gelling within the targeted 5-15 minute window. Specifically, reactions during the 4D characterization campaign were conducted at 30 °C, whereas the bioactivity assays were done without that control. The lower temperature in the bioactivity assays (conducted at room temperature) likely explains the slower gelation observed in those samples. These discrepancies highlight the potential need to characterize a biomaterial system in a manner that anticipates the intended use as closely as possible. On the other hand, the ability to reproduce gel times in two of the four selection treatments in spite of the experimental differences suggests that certain regions of the bacteria-embedded silk formula space are quite robust.

Biological activity in the treatments was assessed according to three criteria that could be observed microscopically. The first criterion was the relative number of cells, as suggested by the number of *E.coli*-sized fluorescent puncta. The second was the amount of fluorescent protein in the cells, as judged by the brightness of the individual puncta. The third was the amount of growth as measured by the morphology of a colony extending beyond individual *E.coli*-sized puncta.

By visual inspection, cell numbers appeared roughly constant across treatments. As each treatment was seeded with a matched bacterial population, the observation of similar quantities of *E.coli*-sized puncta across treatments suggests no significant differences in reproduction. For each of the four treatments, fluorescence intensity in *E.coli*-sized puncta was high near the glass surface and generally diminished as the optical pathlength increased. As the availability of oxygen, required for fluorophore maturation, is expected to be higher as the distance from the coverglass increases, the observed attenuation in the fluorescence intensity with increased distance from the coverglass is attributed to optical attenuation in the material. In fact, the clarity of distinct silk hydrogels does differ depending on formula (data not shown), and the formula with the highest H_2_O_2_ concentration demonstrated the most significant optical attenuation. The variations in optical attenuations made it impossible to compare the treatments by fluorescence intensity differences alone without complex normalization schemes.

Despite the challenges in comparing treatments based on simple fluorescence intensity, **Figure 5** presents evidence that bacterial growth within each of the distinct formulations was qualitatively similar. Near the gel-liquid interface for each formulation, cells grew into sphere-like colonies. The thickness of the region producing the colony formations was similar across treatments. Cells far from the gel-liquid interface did not manifest similar colony morphology. For the two cases which repeatedly gelled within the chosen window of 5-15 minutes, differences in confocal images are not pronounced. This observation suggests that other material properties, for example stiffness and optical clarity, may be able to be tuned independent of the gel time and biological productivity. Further studies on bacterial growth within hydrogels are needed to study the range in which material properties can be varied while maintaining bacterial activity.

## 3. Conclusion

Exciting opportunities exist for living biomaterials. Challenges to their development and perfection include the potential vastness of material formulation spaces and potential trade-offs in competing materials and biological properties. Tools to facilitate rapid screening of large numbers of formula permutations and processing conditions are needed to maximize return on development efforts.

We modified a recently developed tool for microrheology known as differential dynamic microscopy to enable a new framework for automated screening of gel times in bacteria-laden silk hydrogels. The last point in the gelation process for which DDM resolves physically meaningful MSD curves was developed as a simple criterion for the onset of gelation. This approach, termed DDM-RL, is shown to correlate approximately with gel time measured by MPT-*G*′/*G*′′ cross-over in hydrogels comprised of enzymatically crosslinked silk. DDM-RL thus provides a supervision-free route to the approximate determination of gelation kinetics which is valuable for automated or high-throughput screens.

In hydrogels with added *E. coli* cells, a range of gel times is observed when formula components (silk, horseradish peroxidase, hydrogen peroxide, and bacterial cells) are varied over wide ranges. Within a small subset of treatments that gel within 5-15 minutes with matched bacterial loads, biological activity as measured by production of fluorescent protein appeared to be roughly the same. This suggests that other material properties, for example stiffness and optical clarity, may be able to be tuned independent of the gel time while maintaining biological activity.

The mix of properties important to the success of biomaterial design efforts, in combination with the potential vastness of formulation spaces, highlights the need for tools to facilitate rapid and intelligent screening. In this regard, we demonstrated that a combination of Bayesian ML models and optimal decision-making under uncertainty was more efficient than an expert user at choosing experimental treatments to characterize a complex formulation space. Interestingly, the data obtained when switching planner responsibility between an ML algorithm and expert user suggests the possibility that cooperative experimentation between ML and experts could result in increased learning in comparison to either agent acting alone.

## 4. Experimental Section

### Silk hydrogelation

Silk hydrogels are made according to previously published procedures^[22]^ which are based on di-tyrosine crosslinking by horseradish peroxidase (HRP) in the presence of hydrogen peroxide. A two-part mixing scheme is employed to control gelation and mixing by separating the HRP from the hydrogen peroxide until the time of mixing. Part A contains water, silk, cells, and HRP. Part B contains water, silk, cells, and hydrogen peroxide. Tracer beads are added to the silk formulas at 1:700 dilution, resulting in approximately 100-150 beads per 40 x 40 μm^2^ field of view. Beads are near-IR calibration beads (750 nm excitation) with a diameter of 100 nm, dialyzed against water to remove cytotoxic sodium azide. (F8799, Thermofisher).

Silk was isolated from *Bombyx mori* cocoons according to previously published procedures.^[6a, 22]^ In brief, cocoons were obtained commercially (e.g., Aurora Silk, Portland, Oregon, USA), delaminated and scraped clean of debris, boiled in sodium carbonate in water, soaked in lithium bromide in water, and dialyzed. Final silk concentration was assayed gravimetrically. Typical silk yields were approximately 50 milliliters of 6.5% w/v silk from 7.5 grams of dewormed, delaminated cocoons.

Formulation treatments were prepared from frozen stock aliquots. Silk, HRP, and bacterial cell aliquots were all stored at −80 °C at concentrations of roughly 6% silk, 2000 U/ml HRP, and cells at 10x OD_600_ of 1.25. Stock hydrogen peroxide (0.65M) H_2_O_2_ was stored at 4 °C. Prior to use, a silk aliquot was thawed to room temperature and then lightly spun down. In some cases, a miniscule pellet of precipitated silk would appear on the tube side wall, but precipitated silk was not added to experimental samples.

Throughout this work, cell fractions in the silk formulations are expressed as cells/ml, based on a conversion of 8 x 10^8^ cell/ml per OD_600_ unit. Cells were added to silk hydrogels in fractions ranging from ∼0.13 – 1.43 OD_600_ equivalents, corresponding to approximately 0.1 – 1.4 x 10^9^ cells/ml.

### Bacterial culture

*E. coli* BL21 (DE3) was transformed with the pDawn plasmid and used for all experiments involving bacterial viability and induction. pDawn puts expression of dsRed-Express2 under white light illumination control^[32]^ and carries a resistance marker for kanamycin. Bacterial aliquots for mixing into hydrogels were prepared according to two protocols, depending on the experiment. For experiments characterizing living material gel time in 4D formulations (as shown in **Figure 4**), a large number of bacterial aliquots were made of overnight bacteria harboring the pDawn plasmid. Bacteria were first shaken in LB at 37 °C with kanamycin (50 μg/ml) overnight, and then the overnight culture was spun down and resuspended in PBS (pH 7.4). The suspension was then titrated to an OD_600_ of 1.25 using PBS and then pelleted and resuspended in water to 1/10 volume for a 10x concentration of OD_600_ of 1.25. Aliquots of 120 μl were placed into a −80 °C freezer and stored until use, thawed immediately before use and kept on ice during use. All aliquots were used within one week.

For experiments characterizing biological production of bacteria in silk hydrogels (as shown in **Figure 5**), cells were first grown overnight in LB with 50 μg/ml kanamycin. 400 μl of overnight culture was then added to 20 ml M9 minimal medium, supplemented with vitamins, amino acids, and kanamycin again at 50 μg/ml. Cells were grown for 4-5 hours until an OD_600_ of approximately 1. Cells were then spun down and concentrated in M9 salts to achieve an OD_600_ of 1.25. The cells were then pelleted again and resuspended in 1/10 volume water and placed on ice to achieve a concentration matching bacteria used in the gel time experiments. Cells were kept in the dark during preparation of aliquots in order to keep background fluorescent protein to a minimum. Within 20 minutes after concentrating into water, cells were added to hydrogel precursor solutions and mixed together. Gels were cast as 100 μl slabs in 8-well chambered coverglass (Lab-Tek, Rochester NY, USA). Approximately 30 minutes after gel reaction initiation, the samples were overlaid with 600 μl of fresh M9 medium with antibiotics and supplements and placed into an incubator at 37 °C. Cells were grown overnight with continuous, incandescent illumination to induce protein expression (dsRed-Express2) from the pDawn plasmid.

### Differential Dynamic Microscopy Microrheology

Samples ranging in volumes from 100-400 μl were imaged within open (unsealed), 8-well chambered coverglass slides (Lab-Tek, Rochester NY, USA). For experiments characterizing the 4D formulation space, the samples were held at 30 °C throughout the course of gelation using a heated stage whereas replicated treatments for the growth and viability study were at room temperature. Images were taken from a focal plane ∼100 μm above the coverglass. A typical acquisition framerate was 100 Hz, acquired using a CMOS camera on a Vutara 352 microscope (Bruker, Billerica, MA, USA) with a 60x water immersion lens. Tracking bead density was selected to result in approximately 100 beads per frame (per 40 x 40 μm^2^ field of view). Beads were carboxylate-functionalized polystyrene of 100 nm diameter, reporting in the near-IR wavelengths (F8799, Thermofisher). Videos of 1000 frames each were captured at several points as each sample gelled. A typical gelation reaction was observed every 2 minutes over a 30 minute timeframe. Intensity fluctuations in each image stack were converted to an image structure function using DDM implemented in MATLAB, as previously reported.^[25b, 25c, 27]^ The mean-squared displacement (MSD) curves were then extracted from the image structure functions^[25b]^ using a standard deviation acceptance cutoff of 0.025 a.u. in the plateau regions of the *D(q,Δt)* for each wave vector *q*. See **Section S10** in the **Supporting Information** for more detail. Additional convenience code was written as part of this work in order to automate the collection and processing of gelation experiments. See **Section S11** in the **Supporting Information** for details.

### Gel time estimates by MPT and DDM

Gel times are estimated by MPT by computing the MSD at each time point,^[25b]^ using a local power law approximation to compute the *G*′/*G*′′ as functions of frequency,^[33]^ and by identifying the time at which *G*′ exceeds *G*′′ for all frequencies. For DDM-based gel times, estimates were obtained by adding ½ the sampling interval to the time associated with the last MSD curve calculated by DDM. Error in the measurement is reported as ± ½ the sampling interval. As an example, if a gelation reaction was observed every 2 minutes and the last physically reasonable MSD curve reported by DDM occurred at minute 14, then the gel time was estimated at 15 ± 1 minutes. Occasionally, DDM failed to resolve MSDs at some pre-gelation time points but would succeed in later pre-gelation time points. See **Panel B** of **Figure S9** at time points 2.56 and 3.24 minutes for an example. The reason for the failure is not apparent but forced the estimation of gel time by the last time point that DDM resolves an MSD rather than the first time point for which it fails to resolve. See **S12** in the **Supporting Information** for more detail.

In some cases, the last or last few MSD curves of a gelling solution would result in physically nonsensical MSD curves. These curves would appear erratic, relatively independent of lag time, and report MSDs several orders of magnitude higher than earlier (temporally adjacent) MSD curves (See **Supporting Information** section **S13** for an example.) These erratic curves likely result from intensity fluctuations due to background noise or out of plane light which do not correlate with particle motion but nevertheless pass the statistical acceptance criteria.^[25b]^ Erratic curves were discarded manually by inspection.

### Identifying silk formulas promoting maximal biological activity

Bacterial colonies were examined with a Bruker Opterra confocal microscope (Bruker, Billerica, MA, USA), imaging directly through the bulk gels. Image stacks were acquired with an 80 x 80 μm^2^ field of view, 200 μm thick, sampling every 0.5 μm. 561 nm laser illumination was at 1-5% with an integration time of 100 ms. Embedded bacteria were imaged close to the coverglass and extending 200 μm up through the gel and into the cell culture medium. 3D volumes were projected into 2D using maximum intensity projections.

### Machine Learning

After *n* iterations of the campaign loop, the planner builds a GP model *B^n^* = *GP*(*μ^n^*, Σ*^n^*) of the gelation time *f*^*^(*x*), where *x* represents the input parameters of a gelation experiment. Here, *μ^n^*(*x*) and Σ*^n^*(*x*, *x*′) represent the time-*n* mean and covariance functions, respectively, and together, the GP model represents a probability distribution over functions that we believe the unknown gelation time response function *f*^*^ is sampled from, based on the *n* experiments performed. These functions can be defined using standard formulas and a prior belief *B*^0^ = *GP*(*μ*^0^, Σ^0^), which was manually specified as

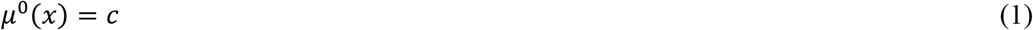

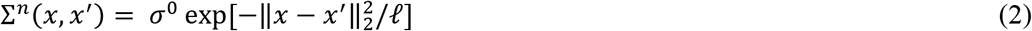

where *c* = 10 s is a prior estimate of the average gelation time we expect to see, *σ*_0_ = 5 s is a prior uncertainty around this initial, constant estimate. The (normalized) length scale *ℓ* = 0.2 describes the presumed statistical relationship between the gelation time *f*^*^(*x*) and *f*^*^(*x*′). See Rasmussen for details^[34]^.

With this belief, we use the Generalized Knowledge Gradient (GKG) decision-making policy to select the next set of experiments to run^[35]^. Namely, we consider the input condition that maximizes

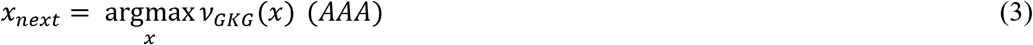

where

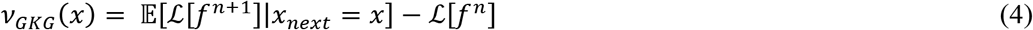

is the GKG acquisition function. Here:

- ℒ[⋅] is a feature operator, that calculates a feature vector from an estimate of the gelation-time response function. Namely, we define a set of query points {*x*_1_, *x*_2_,…, *x_L_*}, and define a bitmap representation predicted optimal region:

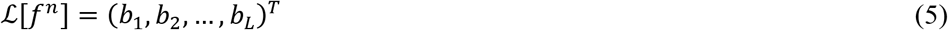

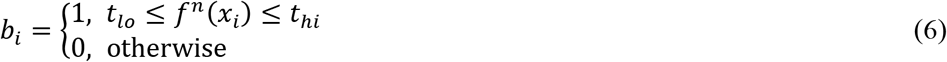
- The expected value term 𝔼[ℒ[*f*^*n*+1^]|*x_next_* = *x*] is the average value of the feature vector ℒ[*f*^*n*+1^] that we would obtain *if* we were to a) run the experiment *x* as the next experiment, b) obtain the output, c) fit the time-(*n* + 1) GP, d) calculate the feature vector from the time -(*n* + 1) GP. It is an average over the result of this *simulation* of what would happen were we to run experiment *x*. In practice, this quantity is calculated using Monte Carlo (MC) methods. An estimate is obtained by averaging over the results of statistical simulations of what would happen were we to run experiment x. The simulated output response is sampled from the time-*n* GP, and subsequent calculations are made upon and averaged over 32 sampled responses.

A four-dimensional grid with logarithmic scaling was used to define query points of the Gaussian process surrogate model. At each grid point *ℓ_i_*, the surrogate was used to define a label *ℓ_i_* to classify it into one of three discrete classes (gelation regimes) based on whether the surrogate predicted a gelation time of less than 5 minutes (*ℓ_i_* = −1), greater than 15 minutes (*ℓ_i_* = 1), or between 5 and 15 minutes (*ℓ_i_* = 0). The grid point was then transformed to a two-dimensional point **z***_i_* = ℎ(*ℓ_i_*) according to the domain-expert defined change in variables. The data set {(*z_i_*, *ℓ_i_*)} was then used to train a kernel Support Vector Machine (SVM) classifier model using a squared-exponential kernel to obtain a classification of points of the 2D space. The colored regions in the predicted gelation regime map indicate the SVM model’s prediction of class at points in the 2D space.

A similar procedure was used to obtain the two-dimensional visualization of model uncertainties shown as the last plot in **Figure 4A**. At each grid point *ℓ_i_*, the GP model was queried to obtain the standard deviation representing the uncertainty *σ_i_* associated with the model’s mean prediction of the gelation time. The data points {(**z***_i_*, *σ_i_*)} were used to train a kernel SVM regressor using a squared-exponential kernel. This SVM model was then used to calculate the projected estimate of the uncertainty in the 2D space.

Error estimates (**Figure 4B**) were computed from the GP model on the gelation time, g(x). Using the hypervolume as metric, the region corresponding to 5 < g(x) < 15 minutes for successive iterates of model fitting (Ω*n*) was compared with the region predicted by the final model (Ω*final*). Relative error was computed as:

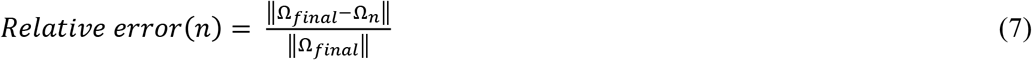

Likewise, the relative change (**Figure 4C**) was computed as follows:

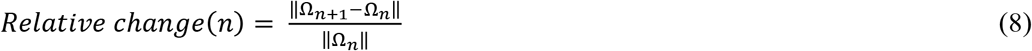

where Ω_*n+1*_ is the predicted hypervolume resulting from several scenarios given a choice of potential experiments.

Note that transformations and the use of SVM models in two-dimensional space were solely for visualization. The decision-making and belief modeling were done in the original 4D space.

## Supporting Information

Supporting Information is available from the Wiley Online Library or from the author.

## Acknowledgements

This work was funded by the Synthetic Biology for Military Environments Applied Research for the Advancement of S&T Priorities program of the Office of the Under Secretary of Defense for Research and Engineering. We thank Jeremy Kemball for assistance with operating the robotic liquid handler.

## References

[1] Y. Zhang, T. Ren, J. He, H. Tian, B. Jin, Colloids and Surfaces A: Physicochemical and Engineering Aspects 2019, 563, 318.

[2] N. Ostrov, M. Jimenez, S. Billerbeck, J. Brisbois, J. Matragrano, A. Ager, V. W. Cornish, Science Advances 2017, 3, 1.

[3] W. Srifa, N. Kosaric, A. Amorin, O. Jadi, Y. Park, S. Mantri, J. Camarena, G. C. Gurtner, M. Porteus, Nat Commun 2020, 11, 2470.

[4] J. Huang, S. Liu, C. Zhang, X. Wang, J. Pu, F. Ba, S. Xue, H. Ye, T. Zhao, K. Li, Y. Wang, J. Zhang, L. Wang, C. Fan, T. K. Lu, C. Zhong, Nat Chem Biol 2019, 15, 34.

[5] P. Q. Nguyen, N. D. Courchesne, A. Duraj-Thatte, P. Praveschotinunt, N. S. Joshi, Adv Mater 2018, 30, e1704847.

[6] a) P. H. Chao, S. Yodmuang, X. Wang, L. Sun, D. L. Kaplan, G. Vunjak-Novakovic, J Biomed Mater Res B Appl Biomater 2010, 95, 84; b) M. Ciocci, I. Cacciotti, D. Seliktar, S. Melino, Int J Biol Macromol 2018, 108, 960; c) O. Hasturk, K. E. Jordan, J. Choi, D. L. Kaplan, Biomaterials 2020, 232, 119720; d) W. He, P. Li, Y. Zhu, M. Liu, X. Huang, H. Qi, New Journal of Chemistry 2019, 43, 2213; e) S. Kapoor, S. C. Kundu, Acta Biomater 2016, 31, 17; f) B. Kundu, N. E. Kurland, S. Bano, C. Patra, F.B. Engel, V. K. Yadavalli, S. C. Kundu, Progress in Polymer Science 2014, 39, 251; g) H. Wu, S. Liu, L. Xiao, X. Dong, Q. Lu, D. L. Kaplan, ACS Appl Mater Interfaces 2016, 8, 17118; h) Z. Zhu, S. Ling, J. Yeo, S. Zhao, L. Tozzi, M. J. Buehler, F. Omenetto, C. Li, D. L. Kaplan, Advanced Functional Materials 2018, 28.

[7] Z. Shao, F. Vollrath, Nature 2002, 418, 741.

[8] a) M. B. Applegate, G. Perotto, D. L. Kaplan, F. G. Omenetto, Biomed Opt Express 2015, 6, 4221; b) N. E. Davis, L. N. Beenken-Rothkopf, A. Mirsoian, N. Kojic, D. L. Kaplan, A. E. Barron, M. J. Fontaine, Biomaterials 2012, 33, 6691; c) S. Yodmuang, S. L. McNamara, A. B. Nover, B. B. Mandal, M. Agarwal, T. A. Kelly, P. H. Chao, C. Hung, D. L. Kaplan, G. Vunjak-Novakovic, Acta Biomater 2015, 11, 27.

[9] A. Baba, S. Matsushita, K. Kitayama, T. Asakura, H. Sezutsu, A. Tanimoto, T. Kanekura, J Biomed Mater Res B Appl Biomater 2019, 107, 97.

[10] J. Chen, H. Venkatesan, J. Hu, Advanced Engineering Materials 2018, 20.

[11] C. Vepari, D. L. Kaplan, Prog Polym Sci 2007, 32, 991.

[12] a) W. Zhang, X. Wang, S. Wang, J. Zhao, L. Xu, C. Zhu, D. Zeng, J. Chen, Z. Zhang, D. L. Kaplan, X. Jiang, Biomaterials 2011, 32, 9415; b) E. Wenk, H. P. Merkle, L. Meinel, J Control Release 2011, 150, 128; c) D. Yu, C. Sun, Z. Zheng, X. Wang, D. Chen, H. Wu, X. Wang, F. Shi, Int J Pharm 2016, 503, 229.

[13] a) X. You, J. J. Pak, Sensors and Actuators B: Chemical 2014, 202, 1357; b) Y. Song, Z. Lin, L. Kong, Y. Xing, N. Lin, Z. Zhang, B.-H. Chen, X.-Y. Liu, Advanced Functional Materials 2017, 27; c) K. Min, S. Kim, S. Kim, Proc Natl Acad Sci U S A 2017, 114, 6185.

[14] R. K. Pal, N. E. Kurland, C. Wang, S. C. Kundu, V. K. Yadavalli, ACS Appl Mater Interfaces 2015, 7, 8809.

[15] a) S. Zhao, Y. Chen, B. P. Partlow, A. S. Golding, P. Tseng, J. Coburn, M. B. Applegate, J. E. Moreau, F. G. Omenetto, D. L. Kaplan, Biomaterials 2016, 93, 60; b) C. J. Bettinger, K. M. Cyr, A. Matsumoto, R. Langer, J. T. Borenstein, D. L. Kaplan, Adv Mater 2007, 19, 2847.

[16] J. Y. Wang, D. Snoeck, S. Van Vlierberghe, W. Verstraete, N. De Belie, Construction and Building Materials 2014, 68, 110.

[17] X. Liu, H. Yuk, S. Lin, G. A. Parada, T. C. Tang, E. Tham, C. de la Fuente-Nunez, T. K. Lu, X. Zhao, Adv Mater 2018, 30.

[18] L. M. Gonzalez, N. Mukhitov, C. A. Voigt, Nat Chem Biol 2020, 16, 126.

[19] S. Sankaran, J. Becker, C. Wittmann, A. Del Campo, Small 2019, 15, e1804717.

[20] M. Schaffner, P. A. Ruhs, F. Coulter, S. Kilcher, A. R. Studart, Science Advances 2017, 3.

[21] S. Mondal, P. Das, A. Kumar Chakraborty, Materials Today: Proceedings 2017, 4, 9833.

[22] B. P. Partlow, C. W. Hanna, J. Rnjak-Kovacina, J. E. Moreau, M. B. Applegate, K. A. Burke, B. Marelli, A. N. Mitropoulos, F. G. Omenetto, D. L. Kaplan, Adv Funct Mater 2014, 24, 4615.

[23] a) J. C. Crocker, D. G. Grier, Journal of Colloid and Interface Science 1996, 179, 298; b) T. G. Mason, Rheologica Acta 2000, 39, 371; c) Y. Gao, M. L. Kilfoil, Optics Express 2009, 17, 4685.

[24] a) V. Breedveld, D. J. Pine, Journal of Materials Science 2003, 38, 4461; b) E. M. Furst, T. M. Squires, Microrheology, Oxford University Press, New York, NY 2017; c) L. L. Josephson, W. J. Galush, E. M. Furst, Biomicrofluidics 2016, 10, 043503; d) K. M. Schultz, A. V. Bayles, A. D. Baldwin, K. L. Kiick, E. M. Furst, Biomacromolecules 2011, 12, 4178.

[25] a) A. V. Bayles, T. M. Squires, M. E. Helgeson, Soft Matter 2016, 12, 2440; b) A. V. Bayles, T. M. Squires, M. E. Helgeson, Rheologica Acta 2017, 56, 863; c) R. Cerbino, V. Trappe, Phys Rev Lett 2008, 100, 188102.

[26] R. Cerbino, D. Piotti, M. Buscaglia, F. Giavazzi, J Phys Condens Matter 2018, 30, 025901.

[27] F. Giavazzi, D. Brogioli, V. Trappe, T. Bellini, R. Cerbino, Phys Rev E Stat Nonlin Soft Matter Phys 2009, 80, 031403.

[28] P. Edera, D. Bergamini, V. Trappe, F. Giavazzi, R. Cerbino, Physical Review Materials 2017, 1.

[29] a) W. Hong, G. Xu, X. Ou, W. Sun, T. Wang, Z. Tong, Soft Matter 2018, 14, 3694; b) T. Larsen, K. Schultz, E. M. Furst, Korea-Australia Rheology Journal 2008, 20, 165; c) T. H. Larsen, E. M. Furst, Phys Rev Lett 2008, 100, 146001; d) T. Moschakis, B. S. Murray, E. Dickinson, J Colloid Interface Sci 2010, 345, 278; e) K. M. Schultz, E. M. Furst, Soft Matter 2012, 8.

[30] J. M. Newby, A. M. Schaefer, P. T. Lee, M. G. Forest, S. K. Lai, Proc Natl Acad Sci U S A 2018, 115, 9026.

[31] a) S. Chen, K.-R. G. Reyes, M. K. Gupta, M. C. McAlpine, W. B. Powell, SIAM/ASA Journal on Uncertainty Quantification 2015, 3, 320; b) R. W. Epps, M. S. Bowen, A. Volk, K. Abdel-Latif, S. Han, K. G. Reyes, A. Amassian, M. Abolhasani, Adv Mater 2020, 32, e2001626; c) A. E. Gongora, B. Xu, W. Perry, C. Okoye, P. Riley, K. G. Reyes, E. F. Morgan, K. A. Brown, Science Advances 2020, 6, 6.

[32] R. Ohlendorf, R. R. Vidavski, A. Eldar, K. Moffat, A. Moglich, J Mol Biol 2012, 416, 534.

[33] M. A. Escobedo-Sanchez, J. P. Segovia-Gutierrez, A. B. Zuccolotto-Bernez, J. Hansen, C. C. Marciniak, K. Sachowsky, F. Platten, S. U. Egelhaaf, Soft Matter 2018, 14, 7016.

[34] C. E. Rasmussen, in Summer School on Machine Learning, Springer, Berlin, Heidelberg 2003.

[35] K. G. Reyes, F. J. Alexander, in Handbook on Big Data and Machine Learning in the Physical Sciences, Vol. 2 (Eds: K. Kleese van Dam, K. G. Yager, S. I. Campbell, R. Farnsworth, M. van Dam) 2020, Ch. 13.

