## Supporting Information for "Engineering Gelation Kinetics in Living Silk Hydrogels by Differential Dynamic Microscopy Microrheology and Machine Learning"

#### S1. Gel time estimation methods

A number of methods for gel time characterization exist. Each of these presents difficulties for automated application. In the simplest case, a vial of gelling material is inverted and monitored for flow. However, surface tension in low volume samples and high viscosity in some other samples can each retard flow even in the absence of gelation. This lack of an unambiguous gelation criterion makes it difficult to determine when gelation has occurred. Further, its disconnect from theoretical foundations yields no additional rheological information even when gelation has been properly assessed. In contrast, bulk rheology techniques including oscillatory rheology can be used to rigorously measure gelation kinetics and other mechanical properties including complex shear modulus and viscosity. However, these traditional techniques are not well-suited to high-throughput or automated application due to lengthy measurement cycles and painstaking sample preparation and setup. They also require relatively large volumes for testing.

Low volume measurements can be made using passive microrheology. Passive microrheology uses the thermally driven motion of probe particles to measure the viscoelastic properties of small volumes of soft materials, especially those in the low stiffness/low viscoelasticity regime.[1] Particle motion is observed using optical microscopy, and multiple particle tracking (MPT) algorithms are then conventionally employed to determine the mean-squared displacement (MSD) of the tracer particles. The MSD encodes the material's viscosity, gel time, and even complex shear modulus by way of the Generalized Stokes-Einstein Relationship (GSER).[1a, 2] Once MSDs are obtained, a number of techniques exist to convert sets of MSDs obtained during a gelation reaction into an estimate of the gel time.

As a low-volume, non-contact, and non-destructive method, passive microrheology using MPT has been successfully used for high-throughput materials characterization.[3] MPT requires relatively simple optical equipment which is commonly available across research laboratories. The major limitation of MPT applied to fully automated microrheology is that the particle localization algorithms require interactive tuning of several processing parameters by an expert operator.[4]

**S2. Estimation of gel time by MPT-G′/G″**

Assuming MSDs are available which capture a suitable range of gelation stages, a number of methods have been developed to extract the gel time. One common approach assumes an MSD curve near the gel point will have a power law exponent of approximately 0.5.[3a] A set of MSD curves can then be examined for power law exponents, and by interpolation or extrapolation the time point that exhibits an exponent of 0.5 can be used to estimate the gel time. This method is only approximate for most gelling materials,[1c] and the range of power law exponents at the critical gel point vary dramatically from as low as 0.11 to up to 0.92.[5]

A more fundamentally justified method for gel time estimation involves calculating the frequency-dependent complex shear modulus $G^*(\omega)$ from the MSDs obtained over the course of a gelation reaction. The complex shear modulus can be used to extract the loss angle, $\delta$, and the time point (or other reaction coordinate) at which the loss angle becomes frequency-independent is the gel time,[1a] Alternatively, the complex shear modulus $G^*$ can be decomposed into its storage ($G'$) and loss components ($G''$) and the time point at which $G'$ is roughly equal to or exceeding $G''$ over all measured frequencies can be used as the gel time.[5] Although this method is attractive as it is based on a fundamental, theoretical understanding of gel point, it is somewhat impractical for automated implementation for a number of reasons. First, estimation of $G^*$ can be unreliable due to instabilities associated with the mathematical transformation of MSD into complex shear modulus by way of inverse Fourier or Laplace transforms.[2a] This is especially true for data of limited frequency content such as generally results from optical microrheology.[2a] In practice, this limitation can be overcome by approximation of $G^*$ and its components $G'$ and $G''$ by a local power law fit of $\alpha(\omega) = |[\delta \ln\langle\Delta r^2(\tau)\rangle/\delta \ln \tau]|_{\tau=1/\omega}$.[2a] However, observing the gelling sample at the exact time at which the loss angle $\delta$ becomes frequency-invariant (or where $G' \sim G''$ over all frequencies) may be difficult to achieve. On the other hand, estimating the time at which the magnitude of $G'$ exceeds the magnitude of $G''$ over all observed frequencies can be used as an approximate method.[5] In this way, one path forward is to collect MSDs during gelation, compute estimates of $G'/G''$ using approximate methods, and then to observe or interpolate the time at which $G'$ exceeds $G''$ over all frequencies.

**S3. Time-cure superposition**

Time-cure superposition (TCS) is both a fundamentally supported and practically accessible method to estimate a material's gel-point[1a]. Assuming a self-similar correspondence between the gelation reaction coordinate (e.g. cure time) and the inverse relaxation time of the material, estimation of gel time is based on assembly of master curves from a family of MSD or modulus curves collected from the gelling sample[6]. Precise time-cure superposition requires convergence of MSD curves in both the pre-gel and post-gel regimes. Even when pre- and post-gel MSD curves are available, the assumption that a given material system will exhibit superposing time-cure behavior may not be universally valid. Appropriate relaxation times for the system need to be sampled in order to capture the features needed for time-cure superposition[5]. Furthermore, assignment of curves to pre- and post-gel regimes is not unambiguous. Assignment of curves is, however, facilitated with complementary knowledge of G′ and G″[1a]. In that case, time-cure superposition essentially reduces to G′/G″ cross-over analysis[7].

**S4. Sensitivity range comparison of DDM and MPT**

Differential dynamic microscopy (DDM) and multiple particle tracking (MPT), although well-correlated for early stage gelation rheological assessment, differ in performance at later stages of gelation. See **Figure S1** below. In the case of silk gelation (in this case, without added bacterial cells), DDM reports MSD curves only for time points of 14 minutes or less. For all these times, the power law exponent exceeds 0.5, a traditional heuristic cutoff for gelation. MPT, on the other hand, continues to report MSD curves for the entire duration of the experiment (30 minutes in this case). The slope of the MSD curves drops from near 1, through the heuristic cutoff of 0.5, and downwards to essentially zero. MSD is thus better able to capture gelation kinetics, and is amenable to the additional gel estimation technique of time cure superposition, as shown at right in **Figure S1** below.

However, the lack of user-selectable parameters for DDM is a compelling advantage over MPT, especially where high-throughput or automated microrheology is desired.

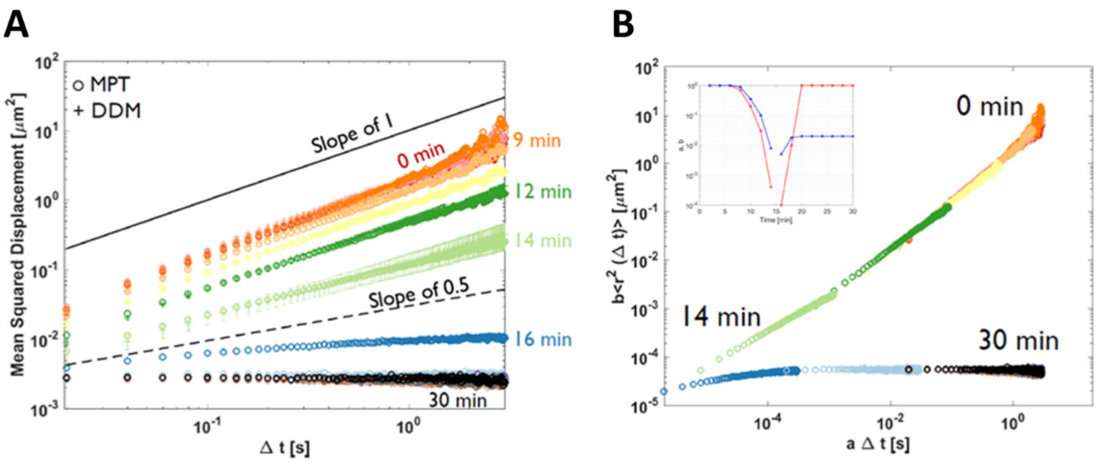


**Figure S1. Sensitivity range comparison of DDM and MPT**. (**A**) A limitation of the DDM technique is a more limited dynamic range in comparison with MPT. This is demonstrated above where a silk hydrogel comprised of 3% silk, 100 ppm $H_2O_2$, and 20 U/ml HRP was reacted under a video microscope and then processed by both DDM and MPT. MPT, with proper processing parameters, traces the gelation reaction at time points along the reaction trajectory sufficient to observe an MSD drop significantly lower than the slope of 0.5. DDM, in contrast, fails to report MSD curves after 14 minutes post-initiation. The slope of the last MSD curve remains above 0.5. (**B**) The MSD curves produced by MPT are amenable to analysis by time-cure superposition, indicating a gel time of approximately 15 minutes.

It is worth noting that a recent enhancement to DDM extends its practical dynamic range by using two cameras capturing at different frame rates, enabling collection of both high-frequency and long lag time data at far-reduced data consumption rates. Known as cross-differential dynamic microscopy (CDDM),[8] it may enable more precise estimations of gel point and could be used to improve upon the method presented herein. CDDM conceivably enables better conversions of the MSD curves into complex shear modulus or viscosity by extending the range of observed frequencies. Alternatively, the higher sampling density afforded by CDDM facilitates sampling a gelling reaction as close as possible to its gel time.

**S5. Comparison of microrheology by DDM and MPT**

MSDs in this work were obtained using both DDM and MPT. The MSDs obtained by MPT were used to produce the G'/G'' plots after applying a dedrifting algorithm[9]. **Figure S2** below compares the MSDs resulting from each processing step along the path to dedrifted G'/G'' plots. The dedrifting algorithm alters the MSDs significantly in a few cases, removing upturns in the MSD curves at later lag times, indicating that drift in the samples was present but correctable. Gel time estimates by DDM and by MPT-G'/G'' (using dedrifted MSDs) was comparable in these cases, as shown below and in the main text.

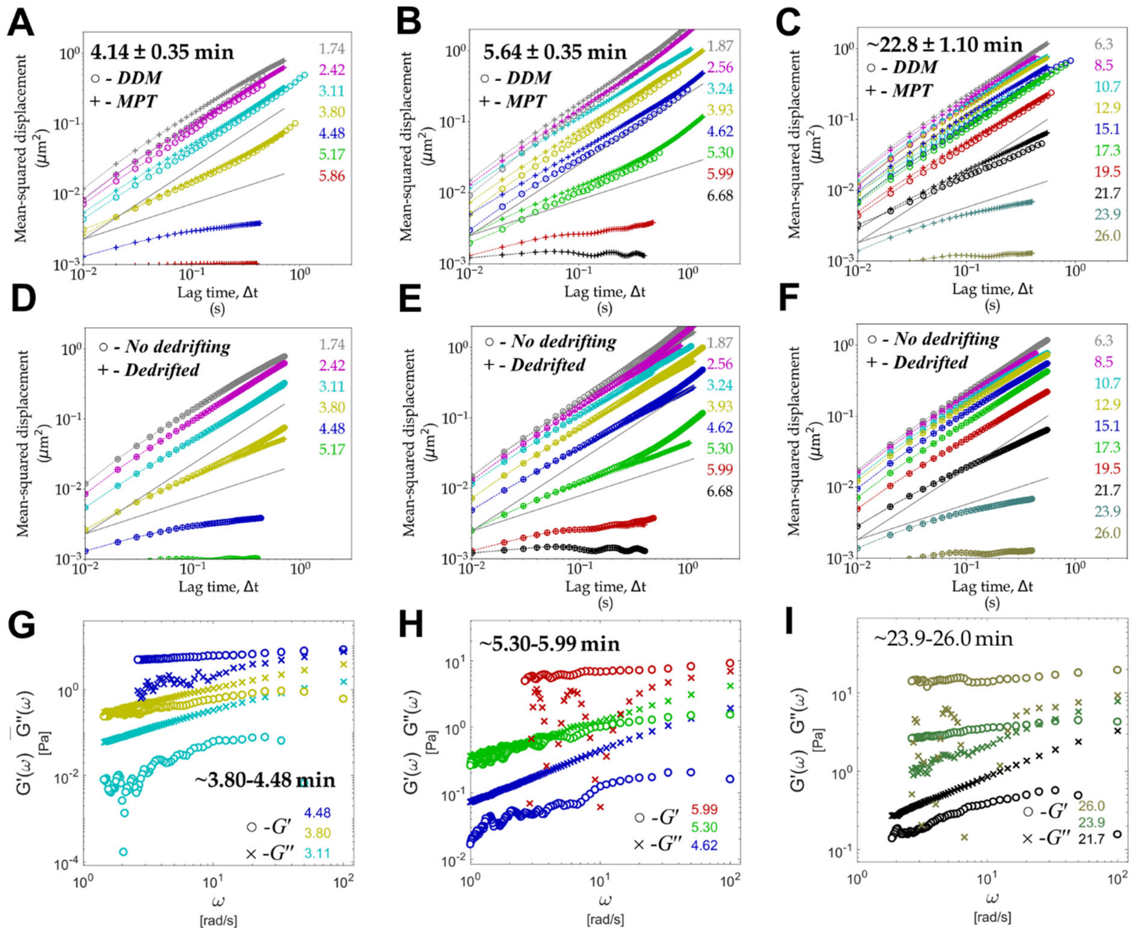


**Figure S2. DDM and MPT microrheology comparison**. Microrheology data is presented for three gelling systems. Each column of plots follows processing for each gel formulation. MSDs obtained by both DDM and MPT are plotted in **Panels A-C**, without dedrifting algorithms applied. DDM fails to resolve MSDs after a certain time point in each case, beyond which MPT still resolves MSDs. This is the basis for the gel time estimation method termed DDM-RL (DDM resolution limit), with estimates shown in each plot in **Panels A-C**. **Panels D-F** compare MDSs obtained by MPT with and without applying a dedrifting algorithm, showing drift in some of the samples. MSDs obtained using the dedrifting algorithm were then used to calculate G’/G’’ as shown in **Panels G-I**. Gel time estimates using MPT-G’/G’’ using dedrifted MSDs annotate **Panels G-I** and are in reasonable agreement with estimates obtained by DDM-RL.

**S6. Artificial Intelligence and Particle Tracking**

Artificial intelligence, in particular a convolutional neural network (CNN), has been deployed for particle detection[10]. In fact, a web-based service (aitracker.net) based on this CNN enables investigators to upload video stacks and to supply some basic information about the image stack (for example, acquisition frame rate) and to receive both location-annotated video stacks as well as files containing particle localization data and mean-squared displacements. No processing parameters are required of investigators using the online service, and the underlying algorithms in the original reports feature only one adjustable parameter: a standard deviation for particle displacements.

A set of image stacks tracking fluorescent microparticles in gelling silk (3% silk, 100 ppm $H_2O_2$, and 40 U/ml HRP, no bacteria) was fed into the online convolutional neural network and compared with DDM results on the same image stack. **Figure S3** shows the comparison, suggesting excellent agreement between the techniques for early-stage gelation conditions but dramatically poor disagreement in later stages. The failure of the convolutional neural network in tracking the DDM curves at later stages may be due to the changing optical characteristics of the maturing silk hydrogels. As the gelation proceeds, the opacity of the silk hydrogel increases, resulting in an increasingly noisy background intensity and reduced contrast with the tracking particles. A more exhaustive training set that incorporates changes in signal to background intensity ratios, and in general noisier images at later stages of the gelation process, is likely to overcome this limitation. Note also that DDM fails to provide MSD data for advanced stages of a gelation reaction, despite proving to be quite useful for optically dense samples[11]. Conventional particle tracking with attentive manual processing remains superior to both DDM and CNN as presently implemented in terms of dynamic range of measurement (see **Section S4** in **Supporting Information**).

However, the fact that DDM operates by analysis of intensity fluctuations in the image stacks—and not by the explicit tracking of embedded probe particles—imparts robustness to DDM analysis[11a] and makes DDM an attractive option as the basis for an automated characterization platform.

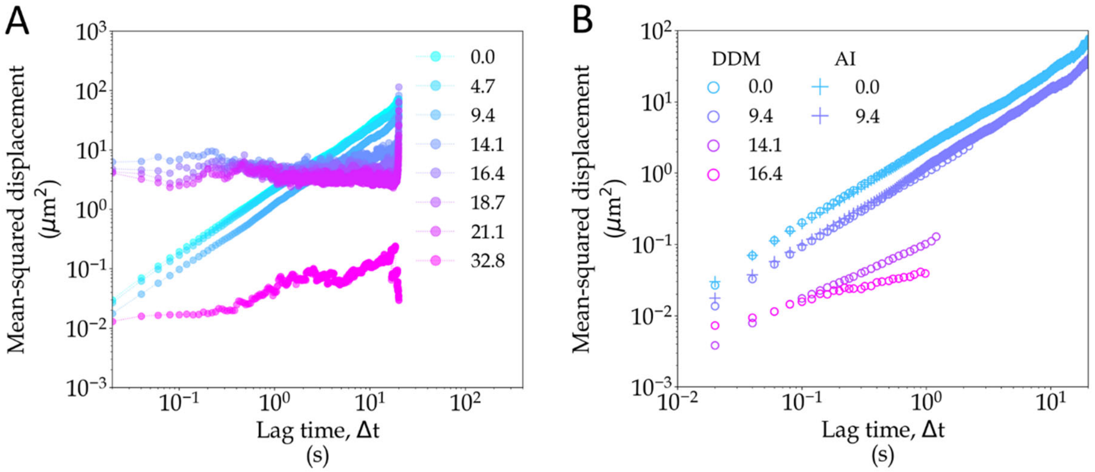


**Figure S3. Comparison of DDM with a convolutional neural network for tracking silk gelation.** Comparison between DDM and CNN tracking[10] is excellent at early stages in the silk gelation course. However, at later points a dramatic divergence occurs. **Panel A** illustrates the MSD calculations of the CNN as a gelation reaction progresses. The first three time points result in smooth MSD curves and capture the changing viscosity of the silk mixture as it gels. However, subsequent curves appear erratic and do not reflect correct rheological properties of the material. **Panel B** illustrates the MSDs as reported by DDM.

Two additional MSD curves are generated, indicating a wider dynamic range in characterizing silk gelation.

### S7. Experimental Planning with Machine Learning

Because MC methods were employed to approximate the GKG-acquisition function values $\nu_{GKG}(x)$, the estimated values obtained are inherently stochastic in nature. See **Figure S4** to see example distributions of these values for three different inputs and five different MC sample sizes. From this, we see that the selected MC sample size of 32, while tightly distributed around its mean and median values, still exhibits a spread of values. Due to this, use of gradient-based optimization to calculate (AAA) is inappropriate. Instead, we perform the optimization using Bayesian optimization. Here, an auxiliary GP model was used to model estimates and uncertainties of the acquisition function, and the expected improvement (EI) decision-making policy was used to query this auxiliary response surface in order to efficiently find the maxima. This global optimization inner loop was iterated 32 times to identify inputs with relatively high acquisition function values.

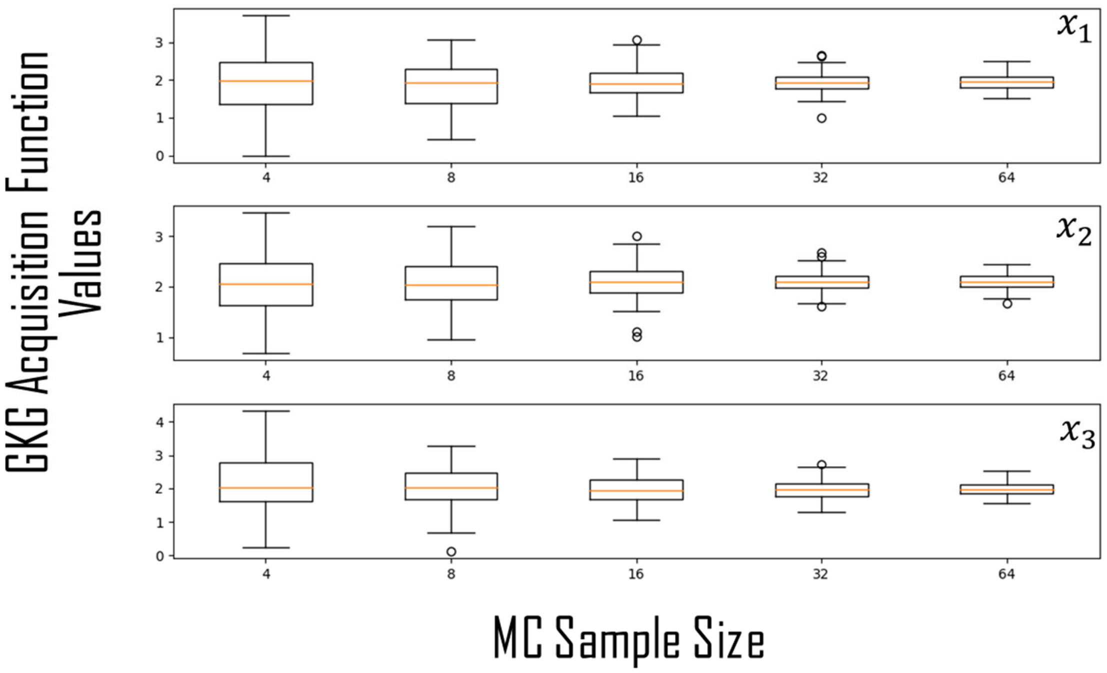


**Figure S4**. The distribution of GKG acquisition function values for different input values (x1, x2, x3) and different MC sample size.

**Table S1. Gel times in bacteria-laden silk hydrogels**

| Treatment | Silk (% w/w) | $H_2O_2$ (ppm) | HRP (U/ml) | Cells ($10^9$/ml) | Gel time (minutes) | $log_{10}$ (HRP*silk/cells) | $log_{10}$ ($H_2O_2$/cells) |
|---|---|---|---|---|---|---|---|
| 1 | 3.13 | 176.00 | 92.00 | 0.76 | 2.5 | 2.58 | 2.36 |
| 2 | 5.56 | 114.16 | 47.14 | 0.62 | 3.5 | 2.63 | 2.27 |
| 3 | 4.85 | 29.22 | 229.42 | 0.94 | 0 | 3.07 | 1.49 |
| 4 | 2.55 | 143.99 | 66.37 | 0.92 | 3.5 | 2.26 | 2.19 |
| 5 | 4.62 | 203.57 | 182.57 | 0.43 | 2.5 | 3.29 | 2.67 |
| 6 | 5.02 | 62.55 | 21.56 | 0.30 | 2.5 | 2.55 | 2.32 |
| 7 | 2.08 | 82.01 | 69.34 | 0.30 | 1.5 | 2.68 | 2.43 |
| 8 | 5.68 | 28.14 | 67.63 | 0.32 | 1.5 | 3.09 | 1.95 |
| 9 | 2.65 | 201.75 | 88.73 | 0.79 | 2.5 | 2.48 | 2.41 |
| 10 | 2.40 | 86.32 | 106.36 | 0.23 | 1.5 | 3.04 | 2.57 |
| 11 | 2.41 | 172.53 | 99.34 | 1.04 | 2.5 | 2.36 | 2.22 |
| 12 | 2.16 | 38.05 | 20.78 | 0.63 | 3.5 | 1.85 | 1.78 |
| 13 | 6.00 | 38.05 | 20.78 | 1.14 | 26.7 | 2.04 | 1.52 |
| 14 | 5.11 | 101.37 | 193.00 | 0.25 | 0 | 3.59 | 2.61 |
| 15 | 4.98 | 29.50 | 218.20 | 0.76 | 0 | 3.16 | 1.59 |
| 16 | 4.50 | 38.05 | 20.78 | 0.86 | 2.5 | 2.04 | 1.65 |
| 17 | 4.89 | 95.19 | 206.58 | 0.88 | 0 | 3.06 | 2.04 |
| 18 | 2.38 | 139.28 | 91.39 | 0.78 | 2.5 | 2.45 | 2.25 |
| 19 | 2.92 | 88.63 | 35.02 | 0.69 | 3.5 | 2.17 | 2.11 |
| 20 | 3.09 | 43.10 | 97.48 | 0.26 | 1.5 | 3.07 | 2.23 |
| 21 | 5.50 | 38.05 | 20.78 | 1.00 | 2.5 | 2.06 | 1.58 |
| 22 | 3.17 | 106.33 | 130.24 | 0.41 | 1.5 | 3.00 | 2.42 |

| Treatment | Silk (% w/w) | $H_2O_2$ (ppm) | HRP (U/ml) | Cells ($10^9$/ml) | Gel time (minutes) | $\log_{10}$ (HRP*silk/cells) | $\log_{10}$ ($H_2O_2$/cells) |
|---|---|---|---|---|---|---|---|
| 23 | 5.35 | 75.00 | 20.78 | 1.14 | 2.5 | 1.99 | 1.82 |
| 24 | 5.35 | 75.00 | 5.00 | 1.14 | 30 | 1.37 | 1.82 |
| 25 | 5.35 | 125.00 | 5.00 | 1.14 | 30 | 1.37 | 2.04 |
| 26 | 5.35 | 150.00 | 7.50 | 1.14 | 30 | 1.55 | 2.12 |
| 27 | 5.31 | 175.00 | 10.00 | 1.14 | 7.5 | 1.67 | 2.19 |
| 28 | 5.31 | 175.00 | 9.00 | 1.14 | 18.4 | 1.62 | 2.19 |
| 29 | 2.00 | 100.00 | 10.00 | 1.14 | 30 | 1.24 | 1.94 |
| 30 | 2.00 | 100.00 | 50.00 | 1.14 | 2.5 | 1.94 | 1.94 |
| 31 | 2.00 | 100.00 | 25.00 | 1.14 | 5.5 | 1.64 | 1.94 |
| 32 | 2.00 | 100.00 | 20.00 | 1.14 | 4.7 | 1.54 | 1.94 |
| 33 | 2.00 | 110.00 | 20.00 | 1.14 | 30 | 1.54 | 1.98 |
| 34 | 2.20 | 125.00 | 23.00 | 1.14 | 20 | 1.65 | 2.04 |
| 35 | 2.50 | 125.00 | 12.00 | 1.14 | 30 | 1.42 | 2.04 |
| 36 | 2.50 | 250.00 | 10.00 | 1.14 | 30 | 1.34 | 2.34 |
| 37 | 3.00 | 300.00 | 25.00 | 1.14 | 13 | 1.82 | 2.42 |
| 38 | 3.00 | 50.00 | 28.00 | 1.14 | 2.2 | 1.87 | 1.64 |
| 39 | 3.00 | 50.00 | 18.00 | 1.14 | 8.12 | 1.67 | 1.64 |
| 40 | 3.00 | 75.00 | 18.00 | 1.14 | 10.6 | 1.67 | 1.82 |
| 41 | 3.30 | 66.31 | 140.11 | 0.45 | 0 | 3.01 | 2.17 |
| 42 | 1.71 | 135.15 | 19.94 | 0.83 | 30 | 1.61 | 2.21 |
| 43 | 2.49 | 84.15 | 103.43 | 0.10 | 1.5 | 3.40 | 2.91 |
| 44 | 4.93 | 43.31 | 176.56 | 1.05 | 0 | 2.92 | 1.61 |
| 45 | 5.25 | 136.54 | 180.54 | 0.44 | 0 | 3.34 | 2.50 |
| 46 | 2.86 | 82.77 | 143.32 | 0.25 | 1.5 | 3.21 | 2.51 |
| 47 | 3.65 | 120.76 | 32.82 | 0.66 | 2.3 | 2.26 | 2.26 |

| Treatment | Silk (% w/w) | $H_2O_2$ (ppm) | HRP (U/ml) | Cells ($10^9$/ml) | Gel time (minutes) | $log_{10}$ (HRP*silk/cells) | $log_{10}$ ($H_2O_2$/cells) |
|---|---|---|---|---|---|---|---|
| 48 | 3.21 | 129.94 | 44.72 | 0.15 | 5.85 | 2.97 | 2.93 |
| 49 | 2.00 | 125.00 | 20.00 | 1.14 | 9.4 | 1.54 | 2.04 |
| 50 | 2.00 | 110.00 | 25.00 | 1.14 | 4.5 | 1.64 | 1.98 |
| 51 | 3.00 | 250.00 | 20.00 | 1.14 | 30 | 1.72 | 2.34 |
| 52 | 3.00 | 300.00 | 28.00 | 1.14 | 5.85 | 1.87 | 2.42 |
| 53 | 3.00 | 150.00 | 28.00 | 1.14 | 4.65 | 1.87 | 2.12 |
| 54 | 3.00 | 50.00 | 23.00 | 1.14 | 4.5 | 1.78 | 1.64 |
| 55 | 3.00 | 50.00 | 21.00 | 1.14 | 3.5 | 1.74 | 1.64 |
| 56 | 3.00 | 75.00 | 21.00 | 1.14 | 3.5 | 1.74 | 1.82 |
| 57 | 3.99 | 56.94 | 49.22 | 0.49 | 1.5 | 2.60 | 2.06 |
| 58 | 4.21 | 67.39 | 203.50 | 1.08 | 0 | 2.90 | 1.80 |
| 59 | 4.24 | 15.34 | 113.19 | 0.87 | 0 | 2.74 | 1.24 |
| 60 | 5.32 | 144.42 | 81.29 | 0.45 | 2.5 | 2.98 | 2.50 |
| 61 | 4.01 | 22.84 | 36.02 | 0.64 | 1.5 | 2.35 | 1.55 |
| 62 | 3.66 | 209.88 | 243.24 | 0.65 | 0 | 3.14 | 2.51 |
| 63 | 3.99 | 159.43 | 121.82 | 0.58 | 1.5 | 2.93 | 2.44 |

### S8. Fluorescence Induction in *E. coli* embedded in silk hydrogels

*E. coli* BL21(DE3) cells containing the plasmid pDawn[12] were cast into silk hydrogels and examined for their capacity for fluorescence induction. For this experiment, all samples were cast into gels comprised of 2.2% silk, 135 ppm $H_2O_2$, 22 U/ml HRP, and 0.42 x $10^9$ cells/ml. Cell-laden gels were allowed to solidify and then were overlayed with M9 culture medium. The cells in the gels were grown at 37 ˚C overnight in an incubator with incandescent illumination. The cultures were not shaken. Fluorescence in the cells, resulting from the production of dsRed-Express2, was strong when cells were both 1) illuminated by white, incandescent light during the overnight culture ('Induced' in **Figure S5** below); and 2) were located near to the cell culture medium-gel interface ('Surface samples'). Cells near the surface presumably had greater access to nutrients and oxygen and were better capable of having their metabolic wastes removed.

Cells that were not induced and cells that were induced but located deep within the hydrogel showed weak fluorescence. The presence of cells in weakly fluorescent or dark regions was confirmed by increasing the laser excitation intensity. Under high illumination intensity, even the fluorescence from leaky expression is apparent. In the chosen samples, uninduced cells near to the surface displayed greater fluorescence than induced cells deep within the hydrogel.

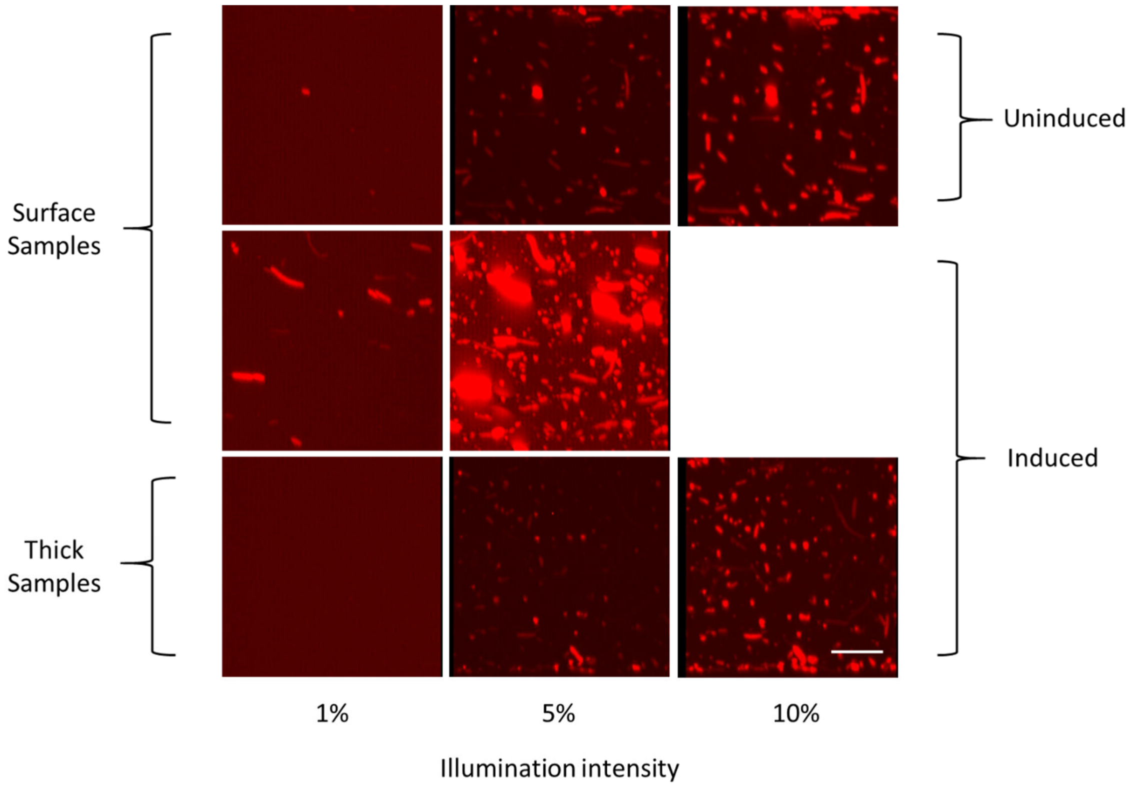


**Figure S5. Fluorescence induction of *E.coli* with pDawn in silk hydrogels**. *E. coli* with pDawn, which produces dsRed-Express2 when exposed to white light, was grown in silk hydrogels with and without illumination. Three samples were examined: 1) cells located far from the gel-culture medium interface grown under incandescent illumination, 2) cells located near to the gel-culture medium interface grown under incandescent illumination, and 3) cells located near to the gel-culture medium interface grown in the dark. Each site was imaged under various excitation intensities ranging from 1-10%. Samples grown near the gel-culture medium interface under illumination produced the strongest fluorescence. Samples grown deep within the gel (far from the gel-culture medium interface) under illumination produced the weakest fluorescence. Cells grown in the dark but near the gel-culture medium interface produced an intermediate amount of fluorescence. Scale bar is 20 μm.

### S9. Colony Morphology in Silk Hydrogels

When cells were inoculated into silk hydrogels at low seeding density, we observed that different formulation conditions promoted differences in bacterial colony formation. That is, replicating cells experienced differences in matrix-derived resistances to proliferation or were otherwise induced into particular growth paradigms by the differences in the gelation processes or components. Presumably, availability of oxygen was not limited for at least some time as growth and production of fluorescence was observed even in thick gel samples. For these experiments, cells were *E. coli* BL21(DE3) constitutively producing the fluorescent protein mEOS3.2[13].

At least three distinct morphologies are produced in response to differences in gelation conditions. These morphologies are depicted in **Figure S6**. Morphologies include oblong spherical, filamentous, and 'hairy' filamentous. These were related to different formula conditions with resultant different gelation kinetics as observed by DDM. Formulas corresponding to the three cases are as follows: 1) oblong spherical morphology (2.65% silk, 17.7 U/ml HRP, 72 ppm $H_2O_2$), gelling in approximately 15 minutes; 2) filamentous (1.85% silk, 18.5 U/ml HRP, 23.1 ppm $H_2O_2$), gelling in approximately 5 minutes; 3) 'hairy' filamentous (1.67% silk, 8.4 U/ml HRP, 21 ppm $H_2O_2$), gel time not measured.

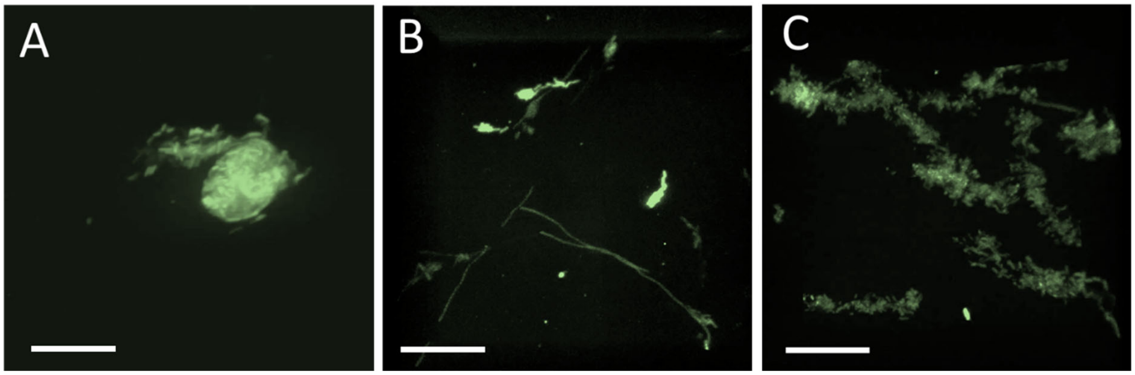


**Figure S6. Colony morphologies in silk hydrogels**. Distinct morphologies of bacterial cells growing in silk hydrogels result from varied silk hydrogel formulations. Oblong spherical (A), filamentous (B), and 'hairy' filamentous (C) can be observed when bacteria are seeded at low density and allowed to grow in gels of different formulas. Scale bars 10 μm.

When perfused with culture medium, cells are able to proliferate and remain viable within silk hydrogels for extended periods of time[14]. Nutrients and waste are able to diffuse in and through the hydrogel matrix. Thus, cells that are capable of producing or sensing molecules of interest can be embedded into the hydrogels to form living biomaterials. These materials can detect and respond to stimuli or can produce and secrete biomolecules.

In certain applications, a high concentration of secreted protein or other biomolecules may need to be produced. In other applications, it may be preferable to maximize the response of each cell within the matrix. For this study, we chose to search for formulation conditions which maximize the average protein concentration within the embedding medium as a measure of the cellular survivability from the gelation process itself. Over the course of the study, however, we inadvertently identified another means by which to tune the functionality of the gel: specifically, the production of bacterial colonies with differing morphology (oblong spherical, fibrillar, or 'hairy' fibrillar). Colonies of varied morphology may present advantages in mass transport for a given application. For example, the delivery of a concentrated bolus of a biological secretion may best be achieved with an oblong spherical morphology, whereas a more evenly dispersed but lower concentration secretion might be delivered by the fibrillar morphology. The potential differences in mass transport may be reflected in the different cellular surface area to cell volume ratios presented in the different morphologies. On the other hand, potential differences may arise due to different metabolic activity of the cells in the various colony morphologies. For example, much of the bacterial biomass in colonies of oblong spherical morphologies may be ineffective at participating in sensing or secretion of biologicals due to poor metabolic conditions of the cells buried within the centers of the relatively thick colonies.

**S10. Extraction of Mean-squared displacements from Image Structure Functions D(q,Δt)**

Production of MSD curves from $D(q, \Delta t)$ data is accomplished according to the general method outlined in Bayles et al[15], but implementation specifics for this study are presented here. In brief, $A(q)$ and $B(q)$ are fit, using the measured $D(q, \Delta t)$, to the expression below:

$$\langle \Delta r^2(\Delta t) \rangle = \frac{4}{q} \ln \left[ \frac{A(q)}{A(q) - D(q, \Delta t) + B(q)} \right] \quad (7)$$

$B(q)$, capturing uncorrelated intensity fluctuations in the measurements, is estimated (for all $q$, such that $B(q) = B$) as the minimum in $D(q, \Delta t_{min})$. $A(q)$, relating to the tracking probe's intensity profile, is estimated as $D(q, \Delta t \to \infty) - B$. This is done by calculating the plateau regions in $D(q, \Delta t)$ as $\Delta t \to \infty$, for each $q$. Identification of appropriate plateau regions and values may vary from implementation to implementation. Parameters that may differ include:

length of plateau region used to estimate *D(q, Δt→∞)*, the standard deviation of *D(q, Δt)* in that plateau region to use as an acceptance criterion, and the location of the plateau region (number of long lag time points to exclude, due to low numbers of observations). For this work, the plateau region was selected as the observations between 70-90% of the Δt indices. That is, the last 10% of Δt observations were ignored for each *q*, and the *A(q)* was calculated as the mean *D(q, Δt)* of the preceding 20% observations. Finally, per Bayles et al[15], lag times where *D(q, Δt)* > 80% of *A(q)* were omitted from $\langle \Delta r^2(\Delta t) \rangle$ calculations, ensuring that the large displacements were not unduly impacted from high *q*, large lag time data. **Figure S7** illustrates how this fitting process is done for one time point in a gelling sample.

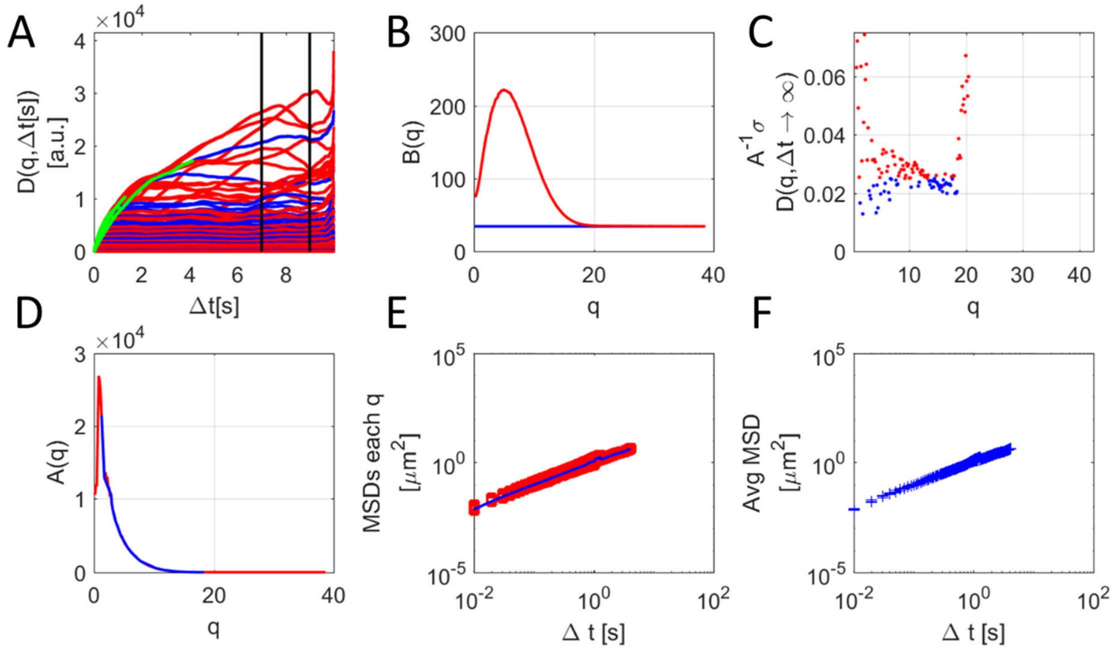


**Figure S7. Deriving MSDs from *D(q, Δt)*.** **Panel A** shows a family of *D* vs *Δt* curves, each curve for a given *q*. Vertical bars indicate the data region examined for plateau behavior. Acceptable curves are indicated in blue, rejected curves in red. The portion of the curves (lag times) with $D(q, \Delta t)$ values less than 80% of $A(q) + B(q)$ are shown in green and used in MSD calculations. **Panel B** shows the estimation of $B(q)$ as the minimum of $D(q, \Delta t_{min})$. $D(q, \Delta t_{min})$ is shown in red, the minimum value shown in blue, plotted as $B(q)$. **Panel C** shows the normalized standard deviations in the plateau region for each $q$, used as rejection criteria for the data in **Panel A**. **Panel D** shows the resultant $A(q)$ vector, with acceptable $q$ colored blue. MSDs for each accepted $q$ are shown in **Panel E** (average MSD shown in blue), and an average MSD with standard deviations from the collection of estimates is shown in **Panel F**.

**S11. Automated Differential Dynamic Microscopy**

Automated application of a MATLAB implementation of DDM (DDMcalc 1.0, https://sites.engineering.ucsb.edu/ ~helgeson/ddm.html) required additional coding. A schematic of the processing pipeline is presented in **Figure S8** below. The source code is available upon request, including documentation for use.

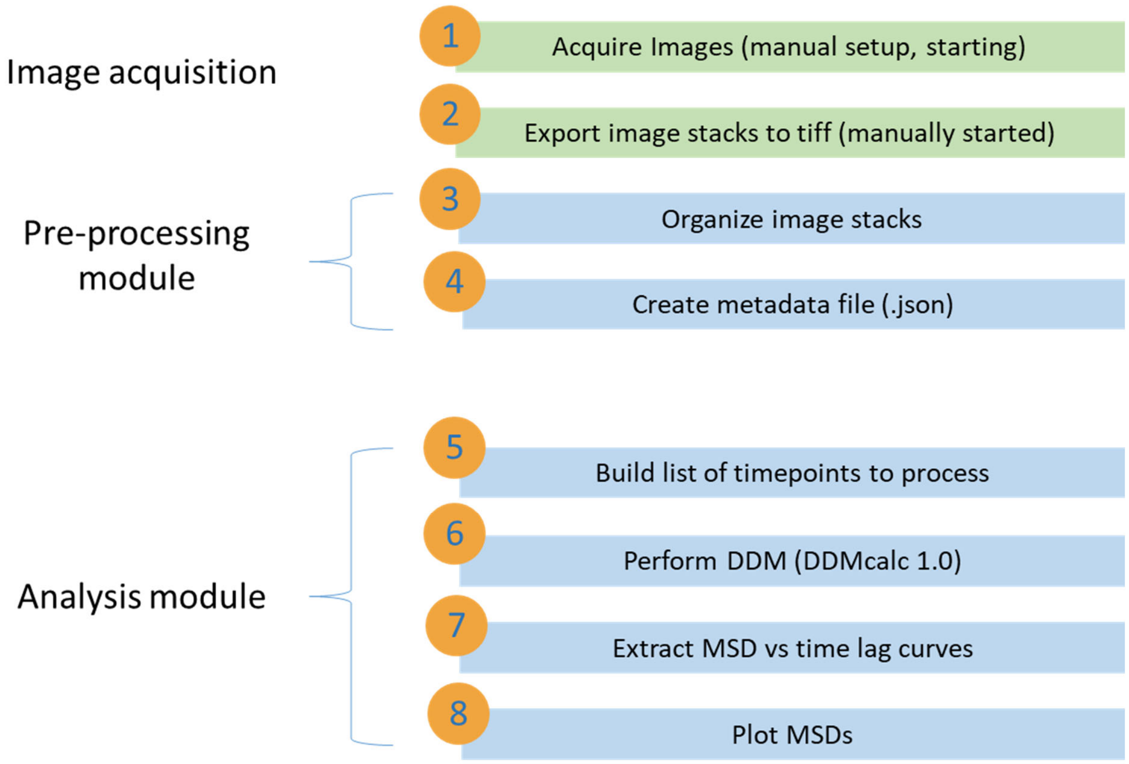


**Figure S8. Automated DDM analysis pipeline**. Two modules enable automated, parameter-free microrheology via DDM starting from raw tiff stacks. The first module normalizes the image stacks, creates an archival folder structure, renames and relocates the image stacks, and creates an experimental parameter sheet (metadata file) which is used to properly process and scale the analyzed data in the second module. The second module creates dynamic structure functions for each image stack for each well and each location in each well, creates an MSD curve for each dynamic structure function, and automatically plots and annotates each series of MSD curves. The gel time for each series of MSD curves is recorded manually and ascertained by inspection.

### S12. DDM failure to resolve due to imaging artefact

Occasionally, artefacts in the imaging will cause DDM to fail. This was observed for the silk hydrogel presented in **Figure S2**, **Panel B** for time points 2.56 and 3.24 minutes. A single frame for each time point is presented below in **Figure S9**, along with a frame exhibiting an imaging artefact that did not result in a failure to resolve (time point at 3.93 minutes). The ability of DDM to resolve the MSD for the later time point, despite the significant optical anomaly, may be due to the more highly restricted motion of the anomaly due to the more advanced stage of gelation. Alternatively, the apparent saturation of the imaging pixels by the extremely bright fluorescence of the probe at time points 2.56 and 3.24 minutes produces non-Gaussian intensity profiles which potentially cause difficulty for fitting A(q) and/or B(q). In contrast, pixel saturation does not appear to be taking place for the anomaly at 3.93 minutes.

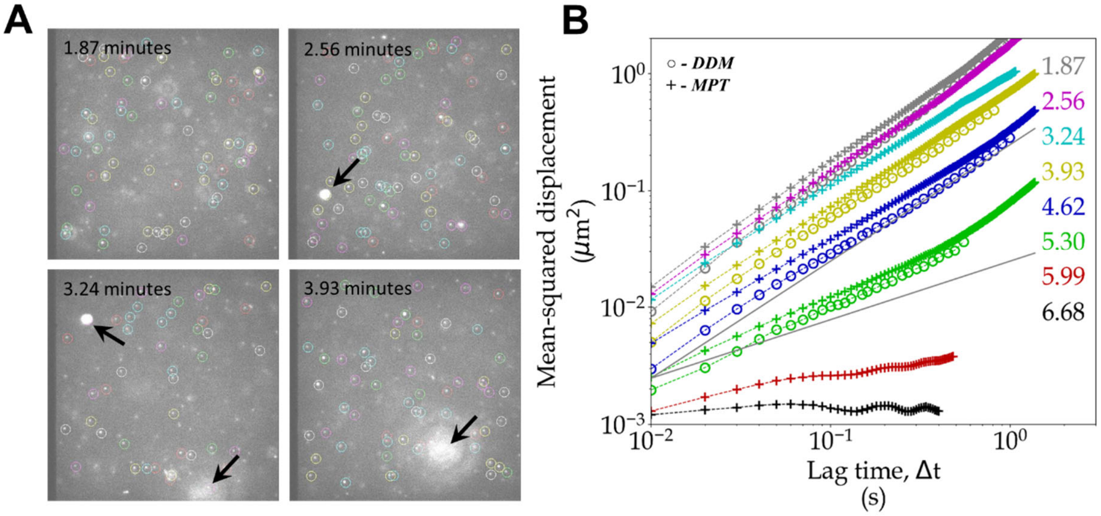


**Figure S9. Imaging artefact interference with DDM**. Imaging abnormalities can impede successful DDM analysis. Frames from videos taken at various time points during a silk hydrogelation are shown in **Panel A**. **Panel B** repeats the DDM/MPT MSDs associated with this experiment. Exceptionally bright spots were observed at 2.56 and 3.24 minutes which prevented successful resolution of MSDs from DDM. An even larger bright spot was observed for later time points (3.93 minutes shown above) which did not prevent resolution of MSDs. The particles at 2.56 and 3.24 minutes, at earlier stages in the gelation, were much more dynamic than the relatively fixed anomoly observed at 3.93 minutes (and beyond). Colored circles in the images indicate localized particles according to MPT. Arrows point to imaging artefacts, presumably associated with exceptionally bright tracer beads or bead aggregation.

### S13. Erratic MSDs

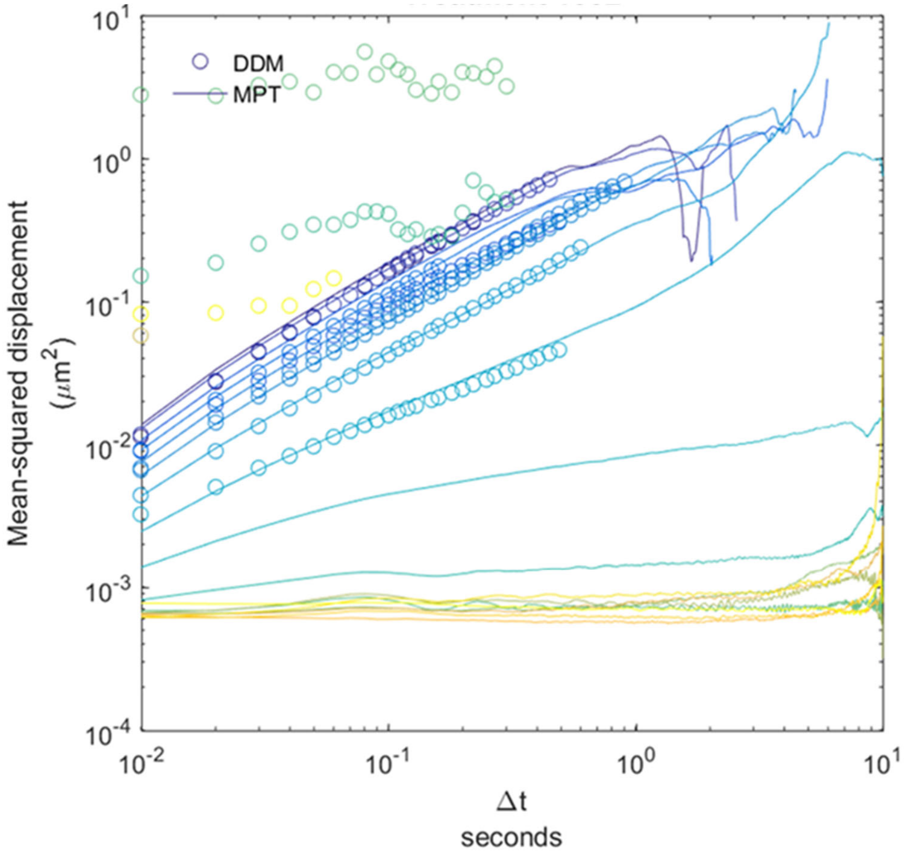


**Figure S10. Erratic MSD from DDM of sample with uncorrelated particle displacements**. In relatively rare cases, the algorithm converting DDM dynamic structure functions into mean-squared displacements yields erratic curves. These curves meet the standard deviation cutoff criterion of the image structure function at a particular time and $q$ (wave-vector parameter). However, it is expected that the intensity fluctuations producing the image structure function at these timepoints is driven by optical scatter in opaque materials, out of plane light, or other sources which do not derive from particle motion. Note the similarity in the results in this failure mode with the results at advanced stages of MSD production by CNN in **Figure S3**.

### References


[1] a) W. Hong, G. Xu, X. Ou, W. Sun, T. Wang, Z. Tong, *Soft Matter* **2018**, 14, 3694; b) T. Larsen, K. Schultz, E. M. Furst, *Korea-Australia Rheology Journal* **2008**, 20, 165; c) T. Moschakis, B. S. Murray, E. Dickinson, *J Colloid Interface Sci* **2010**, 345, 278; d) R. S. Tu, V. Breedveld, *Phys Rev E Stat Nonlin Soft Matter Phys* **2005**, 72, 041914; e) M. A. Woldeyes, L. L. Josephson, D. L. Leiske, W. J. Galush, C. J. Roberts, E. M. Furst, *Mol Pharm* **2018**, 15, 4745.

[2] a) T. G. Mason, *Rheologica Acta* **2000**, 39, 371; b) T. M. Squires, T. G. Mason, *Annual Review of Fluid Mechanics* **2010**, 42, 413.

[3] a) K. M. Schultz, A. D. Baldwin, K. L. Kiick, E. M. Furst, *Soft Matter* **2009**, 5, 740; b) K. M. Schultz, A. V. Bayles, A. D. Baldwin, K. L. Kiick, E. M. Furst, *Biomacromolecules* **2011**, 12, 4178.

[4] J. C. Crocker, D. G. Grier, *Journal of Colloid and Interface Science* **1996**, 179, 298.

[5] E. M. Furst, T. M. Squires, *Microrheology*, Oxford University Press, New York, NY **2017**.

[6] D. Adolf, J. E. Martin, *Macromolecules* **1990**, 23, 3700.

[7] H. H. Winter, F. Chambon, *Journal of Rheology* **1986**, 30, 367.

[8] M. Arko, A. Petelin, *Soft Matter* **2019**, 15, 2791.

[9] Y. Gao, M. L. Kilfoil, *Optics Express* **2009**, 17, 4685.

[10] J. M. Newby, A. M. Schaefer, P. T. Lee, M. G. Forest, S. K. Lai, *Proc Natl Acad Sci U S A* **2018**, 115, 9026.

[11] a) A. V. Bayles, T. M. Squires, M. E. Helgeson, *Soft Matter* **2016**, 12, 2440; b) K. He, M. Spannuth, J. C. Conrad, R. Krishnamoorti, *Soft Matter* **2012**, 8; c) Y. Gao, J. Kim, M. E. Helgeson, *Soft Matter* **2015**, 11, 6360.

[12] R. Ohlendorf, R. R. Vidavski, A. Eldar, K. Moffat, A. Moglich, *J Mol Biol* **2012**, 416, 534.

[13] M. Zhang, H. Chang, Y. Zhang, J. Yu, L. Wu, W. Ji, J. Chen, B. Liu, J. Lu, Y. Liu, J. Zhang, P. Xu, T. Xu, *Nature Methods* **2012**, 9, 727.

[14] I. Drachuk, S. Harbaugh, R. Geryak, D. L. Kaplan, V. V. Tsukruk, N. Kelley-Loughnane, *ACS Biomaterials Science & Engineering* **2017**, 3, 2278.

[15] A. V. Bayles, T. M. Squires, M. E. Helgeson, *Rheologica Acta* **2017**, 56, 863.
